# FERAL: a supervised video-understanding system for direct video-to-behavior mapping

**DOI:** 10.1101/2025.11.16.688666

**Authors:** Peter Skovorodnikov, Jacopo Razzauti, Blair R. Costelloe, Friederike Buck, Vikram Chandra, Dominic D. Frank, Tomas Kay, Benjamin Koger, Orli Snir, Janet Zhao, Iain D. Couzin, Daniel J.C. Kronauer, Leslie B. Vosshall

## Abstract

Quantifying animal behavior often requires segmenting continuous actions into discrete, interpretable states, yet most automated pipelines infer actions from keypoint dynamics and are limited by keypoint tracking quality. Here we present FERAL (Feature Extraction for Recognition of Animal Locomotion), a supervised video-understanding toolkit that maps raw video directly to frame-level behavioral labels, bypassing the keypoint-extraction and pose-classification stages of conventional pipelines. Across benchmarks, FERAL matches or exceeds state-of-the-art pose- and video-based baselines. On the CalMS21 mouse social-interaction benchmark, it exceeds Google’s VideoPrism using only a quarter of the training data. FERAL generalizes across species (apes, zebras, mice, ants, flies, and nematodes), recording conditions, and levels of organization, from single animals to social interactions and colony-scale collective behavior. Released as a user-friendly, open-source package, it integrates with existing analysis pipelines. By enabling scalable, species-agnostic behavioral quantification directly from raw video, FERAL broadens the experimental paradigms available for studying animal behavior.

## INTRODUCTION

Animal behavior unfolds continuously in time, yet is often parsed into discrete, human-interpreted actions [1–4]. Since the earliest days of ethology, researchers have identified these actions through naturalistic observation and manual annotation [5, 6], establishing the foundation of modern behavioral science [3, 7, 8]. Such analyses have been essential across disciplines, informing research in ecology [9], neuroscience [10], neurodegenerative disease modeling, and drug discovery for disorders with observable behavioral phenotypes.

The advent of videotaping in the mid-1980s enabled the quantification of behavior with greater temporal resolution than possible by direct observation alone. Manual annotation remains the most versatile method for quantifying recorded behavior, adapting to scene complexities that range from controlled arenas to field recordings. However, manual annotation is labor-intensive and subjective, and remains the main bottleneck in scaling behavioral research. This burden can be alleviated either by reducing the per-frame labeling cost or by shifting it to the training stage of a classifier that generalizes across recordings [11–13]. Advances in computer vision have partially mitigated this limitation through markerless tracking tools such as DeepLabCut [14], SLEAP [15], and LightningPose [16], which estimate the positions of animals and their body parts over time [17, 18] without the need to label keypoints on the animal manually. These tools have transformed behavioral analysis in laboratory settings and allow scientists to reliably and continuously analyze fine kinematics that are often required to understand the neural basis of behaviors. However, these keypoint-detection tools often require tightly controlled imaging conditions and an experimental setup that minimizes obstructions of such keypoints. These approaches can fail in more naturalistic and noisy environments, forcing researchers to simplify behavioral assays (e.g., by tethering animals, reducing their movements across planes, conducting experiments in a featureless environment with uniform backgrounds, or designing experiment-specific arenas), rely on manual annotation, or abandon experiments involving complex environments or free movement altogether. Importantly, these methods track where animals (and their body parts) are, but not what actions they are performing.

Automatically determining the actions animals are engaged in remains a greater challenge. Pose-based pipelines have advanced two complementary approaches: unsupervised discovery of behavioral repertoires from kinematic motifs (keypoint-MoSeq [19], B-SOiD [20], VAME [21]) and supervised recognition of pre-defined categories (MARS [11], JAABA [22], SimBA [23]). The former is ideal for identifying novel structure in behavioral dynamics where such behaviors depend on the actual pose and spatial organization of animals and their body parts, while the latter is ideal for scaling up the temporal segmentation of specific behaviors when keypoint dynamics are sufficient for reliable detection.

The performance of these approaches, however, depends on the confidence of keypoint detection, and the skeletonized representations they require discard contextual information that can be essential for accurate behavioral interpretation. As the complexity of behavioral organization increases, such as in social or collective dynamics, pose-based pipelines become increasingly demanding, requiring multi-animal tracking and extensive post-tracking curation (**Figure 1a**). In many cases, the visual cues that define a behavior are well established, and researchers primarily need a scalable, automated way to recognize them across large datasets. DeepEthogram [24] was among the first supervised deep-learning toolkits to produce frame-level behavioral sequences directly from raw pixels rather than pose. More recent video-foundation-model approaches — DAP [25], ChimpVLM [26], and PriVi [27] — have pushed this pixel-input direction further, but all three are developed and evaluated on primates and have not been shown to generalize across species. Other efforts have applied video foundation models more broadly to animal behavior [28–30], and recent benchmarks have begun to evaluate video models across scientific domains [31]. However, these remain largely analyses or benchmarks rather than deployable tools: to date, no pixel-input pipeline has been released as a general-purpose, species-agnostic behavioral toolkit spanning recording modalities (e.g., lab, drone, camera-trap) and levels of behavioral organization (e.g., single-animal, pair, collective).

**Figure 1:**
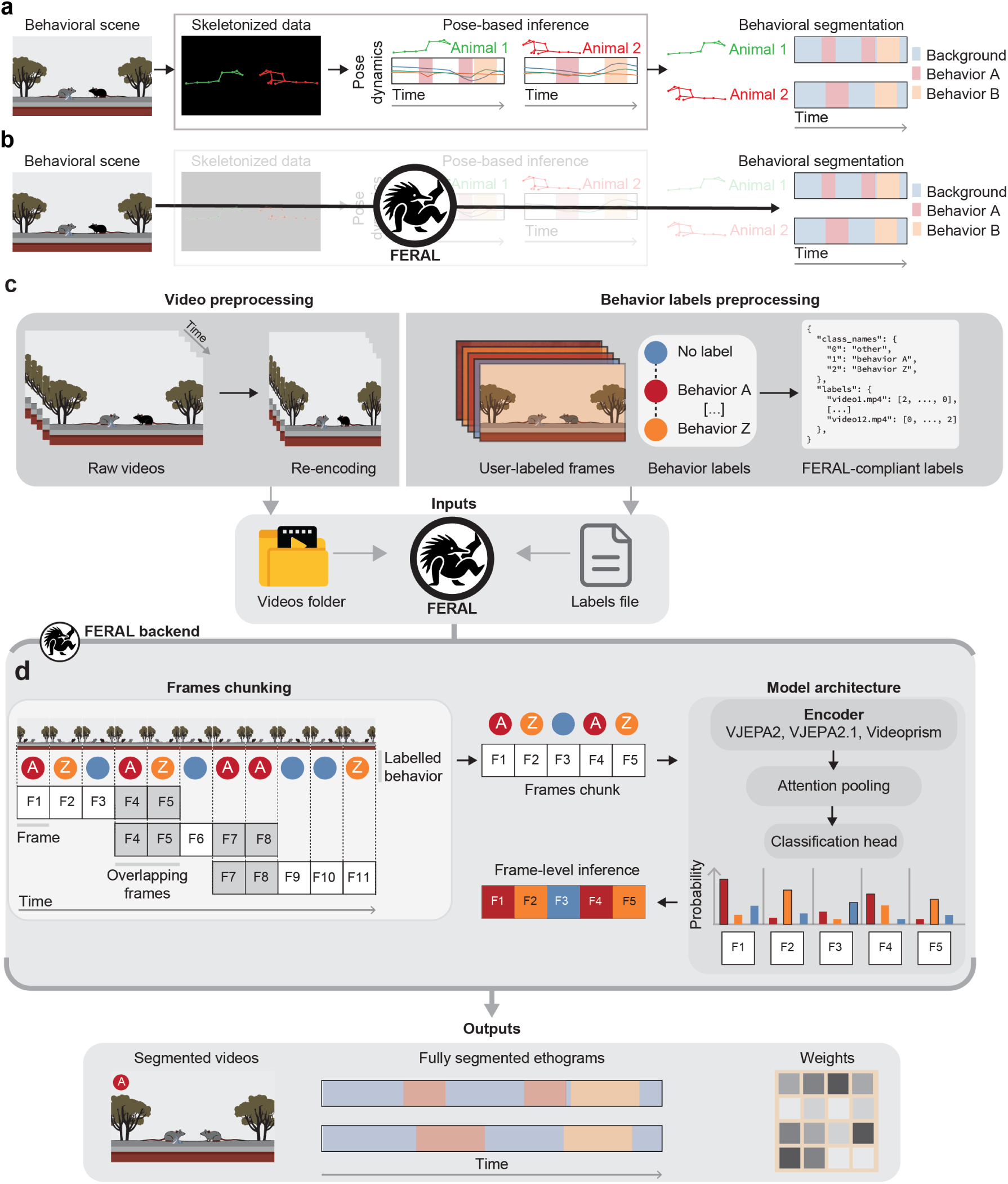
Overview of the FERAL workflow. **(a–b)** Two routes from raw video to behavioral segmentation: the conventional pose-based pipeline (a) versus the FERAL approach (b). **(c)** *Preprocessing*: raw videos are resized and re-encoded into standardized, seekable inputs. User annotations are converted to a common frame-aligned JSON schema (class names and per-frame label arrays). **(d)** *FERAL backend*: videos are divided into overlapping temporal segments. Chunks are then embedded by a pretrained video encoder (e.g., V-JEPA 2); attention pooling aggregates spatiotemporal features, and a lightweight classification head is jointly fine-tuned on the user-supplied frame-level labels. Overlapping predictions are ensembled to produce stable frame-level probabilities.

To bridge this gap, we developed FERAL (Feature Extraction for Recognition of Animal Locomotion), a supervised video-understanding system that learns behavior directly from raw video frames, bypassing the need for pose estimation while preserving full temporal and visual information (**Figure 1b**). This approach is particularly advantageous when pose estimation is unreliable or unnecessary for the desired analysis. FERAL can operate either alongside or independently of existing pose-based frameworks, providing a single, integrated tool for behavioral segmentation.

Leveraging a pretrained video foundation model [32, 33], FERAL integrates motion and visual cues within a unified architecture. Trained directly on videos aligned with user-provided annotations, it detects discrete actions and outputs frame-level behavioral time-series (i.e., segmentations over a user-supplied ethogram, the list of behavior categories of interest) that reliably capture discrete animal behavior.

On benchmarking datasets, FERAL matches or exceeds state-of-the-art pose- and video-based baselines under matched evaluation protocols. It generalizes robustly across datasets spanning diverse species, recording modalities (from controlled laboratory conditions to field and aerial footage), and levels of behavioral organization: from single-animal locomotion in the nematode *Caenorhabditis elegans* (hereafter *C. elegans*) and the fly *Drosophila melanogaster* to social interactions and collective behavior in rodents, ants, and apes. These results demonstrate that FERAL maintains high performance even in scenarios where traditional pipelines typically struggle. For instance, FERAL makes analysis of behaviors involving interaction of animals either with each other or with their environments much more feasible.

Designed for accessibility, FERAL combines preprocessing, training, and inference within a modular workflow requiring minimal coding expertise and at a cost accessible to most researchers. With only a few commands, it can be run locally or on cloud platforms such as Google Colab, providing a scalable, context-aware foundation for modern behavioral science.

## RESULTS

### Direct video-to-behavior mapping with FERAL

FERAL provides a user-friendly workflow that takes raw videos and behavioral annotations as inputs, fine-tunes an open-source video-understanding model, and outputs frame-level behavioral labels (ethograms). To ensure reproducibility across laboratories and recording setups, the pipeline standardizes both inputs (videos and labels) into a common format.

The first stage is *video preprocessing*. Because video-understanding models operate on relatively low-resolution inputs (e.g., by default 256 × 256 pixels for the V-JEPA 2 backbone), input videos must be resized and re-encoded into seekable formats to enable efficient frame sampling [32, 34]. Within FERAL, this feature operates on Linux and Windows operating systems to automate this process, producing standardized inputs regardless of the original recording format (**Figure 1c**).

The second stage, *label preparation*, converts behavioral annotations from diverse software (e.g., BORIS [35], EthoVision [36]) into a consistent schema [35, 37]. The annotations provided by the user, often timestamp-based, are converted into JSON files that map every video frame to one (single class) or more (multiclass) categorical behavioral labels. In practice, users need only a folder of videos and a corresponding label file to initiate training and inference (**Figure 1c**).

Because full-length video recordings cannot be processed in a single pass due to computational limitations of the backbone model, FERAL divides each video into overlapping temporal chunks that are independently processed, and corresponding predictions are subsequently ensembled (**Figure 1d**). This design minimizes misclassification at behavioral boundaries and maintains temporal continuity by ensuring that each short behavior is fully captured within at least one chunk.

After evaluating several potential backbones, we adopted V-JEPA 2 for the FERAL default configuration [32]. V-JEPA 2 is a foundation model from Meta Fundamental AI Research (see Methods). Pretrained on over one million hours of internet video with a self-supervised masked-prediction objective, V-JEPA 2 yields spatiotemporal embeddings that transfer to downstream video-understanding tasks. Within FERAL, we fine-tune the upper twelve of twenty-four transformer layers using user-supplied videos and labels, aligning pretrained representations with the specific demands of behavioral segmentation rather than retraining from scratch [38] (**Figure 1d**).

To generate frame-level predictions, FERAL extends the encoder with an attention-based pooling module and a classification head. Each video chunk is represented as a sequence of spatiotemporal tokens that are integrated by a transformer, then compressed via attention pooling into temporally aligned embeddings. These embeddings are normalized and linearly projected into class logits, producing frame-wise probabilities that are ensembled across overlapping segments.

The resulting outputs are interpretable frame-level behavioral time-series that describe what actions animals are performing at each frame. These can be visualized directly or integrated into downstream analyses. Model weights are automatically saved, allowing users to reuse or fine-tune trained networks on new datasets without retraining from scratch (**Figure 1d**).

### FERAL outperforms state-of-the-art methods across benchmarks

To evaluate FERAL’s precision and robustness relative to existing approaches, we benchmarked its performance on established datasets that provide raw videos aligned with frame-level behavioral annotations. The first, the Caltech Mouse Social Interactions (CalMS21) dataset [39], contains recordings of freely behaving mice (*Mus musculus*) engaged in resident–intruder assays, paired with both tracked poses and expert behavioral labels (**Figure 2a**).

**Figure 2:**
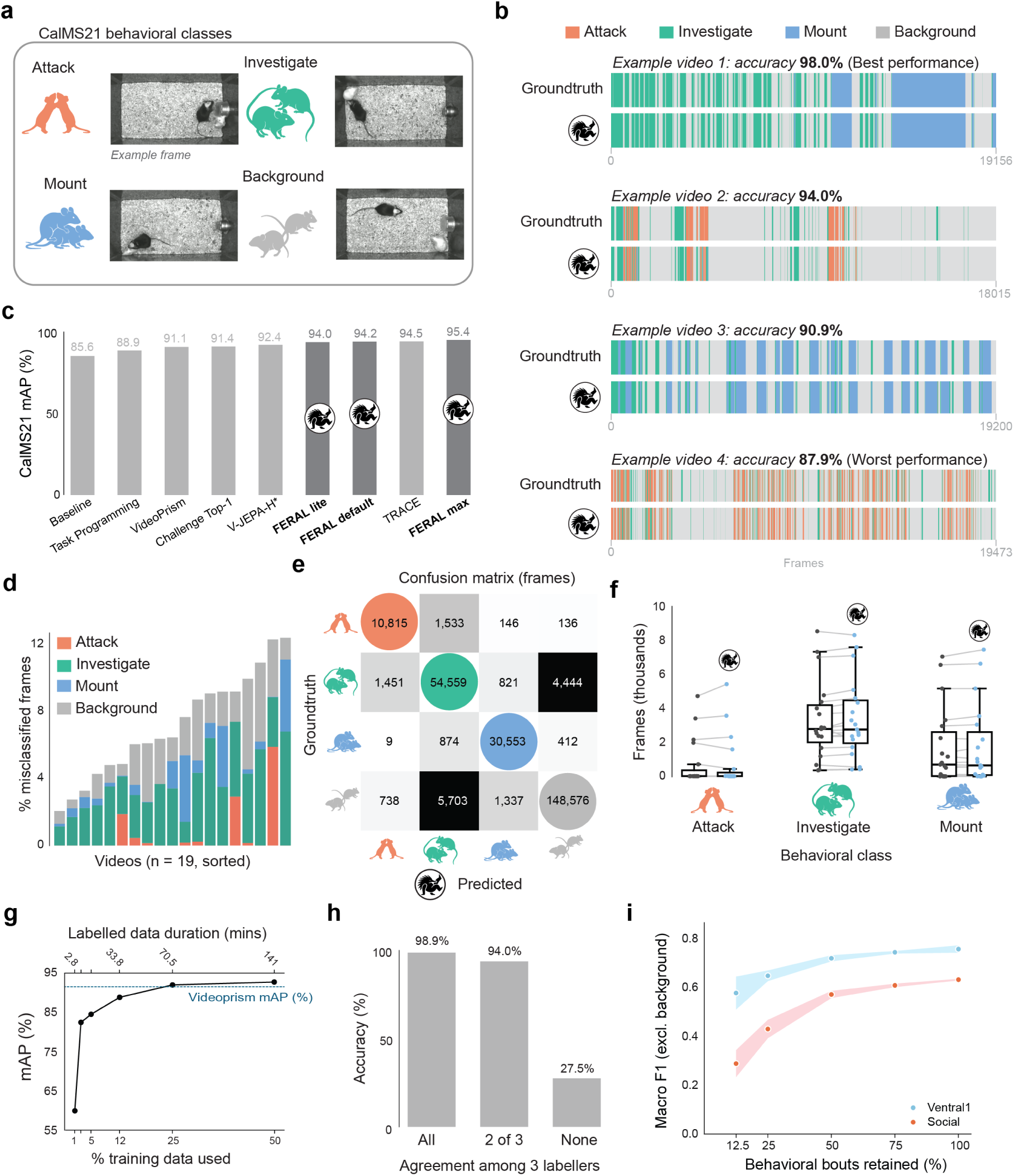
FERAL matches or exceeds state-of-the-art baselines across benchmarks. **(a)** Example frames and corresponding behavioral classes from the CalMS21 dataset. **(b)** Representative frame-level behavioral time-series comparing FERAL predictions (bottom) to ground truth (top). **(c)** Mean average precision (mAP) across models on the CalMS21 dataset including 3 FERAL configurations. **(d)** Fraction of per-class mismatched frames across all CalMS21 videos. **(e)** Confusion matrix of frame-level predictions versus ground truth. **(f)** Comparison between ground-truth and predicted frame counts for each behavioral category across videos. **(g)** Data-efficiency analysis showing mean average precision (mAP) as a function of the percentage of training data. VideoPrism’s performance is taken from the original paper. **(h)** FERAL performance on frames binned by annotator agreement. **(i)** Data-efficiency analysis showing Macro F1 as a function of used behavioral bouts for training.

Originally introduced as a community challenge for the classification of social behavior, the CalMS21 dataset included multiple baseline models and public submissions. We compared FERAL against a suite of alternative pose- and video-based approaches using mean average precision (mAP), the official metric of the original challenge. A random classifier would get 13.5% mAP while a perfect one would achieve 100% mAP. The strongest released baseline, in addition to the labeled behavioral dataset, employed self-supervised pretraining on large sets of unlabeled pose trajectories [39]. We also report the Competition Top-1 entry from the official leaderboard, results from Google’s VideoPrism paper [33], Task Programming [40], VJEPA-H [31] and TRACE [41]. To accommodate compute and runtime limitations of more users and make our tool more broadly accessible, we are providing and benchmarking three distinct configurations for FERAL: *lite*, *default*, and *max*. We discuss GPU VRAM requirements, runtimes, and performance for each of these configurations in the Methods section.

FERAL default reached 94.2% mAP on CalMS21 (**Figure 2b-c, Supplementary Video 1**), exceeding the Sun et al. 2021 [39] self-supervised pose-pretraining baseline, the MABe 2021 challenge Top-1, and results from the VideoPrism paper [33]; all the numbers are mAP under the original CalMS21 challenge protocol. To isolate the contribution of our fine-tuning protocol from that of the backbone itself, we additionally evaluated FERAL using the VideoPrism, V-JEPA 2 and V-JEPA 2.1 backbones and present results in **Table 6**.

FERAL also maintained consistent performance across all test CalMS21 mouse videos, correctly classifying the majority of the frames within each sequence (**Figure 2d**). Confusion-matrix analysis showed that residual errors were mostly confined to confusion between “background” and “investigate” categories, indicating occasional uncertainty in distinguishing background periods from subtle actions (**Figure 2e**). Quantitative comparison of predicted and annotated total behavior durations per video confirmed high agreement across all labeled behavioral categories, demonstrating that FERAL accurately preserved both the temporal structure and balance between classes of the annotated behavioral data (**Figure 2f**).

To assess how much labeled data FERAL requires to achieve strong performance, we progressively subsampled the training set of CalMS21 at the chunk level (i.e., randomly retaining a fraction of training chunks; see Methods, Data efficiency). FERAL achieved a mAP of 92.8% using only 50% of the available training data and 92.1% with 25%, already surpassing both the VideoPrism and Competition Top-1 models trained on the full dataset. Even when trained with only 12% of the data, performance remained high (89.0%), and robust mAP was maintained down to 5% (84.6%) and 2.5% (82.6%). At the lowest sampling level (1%), FERAL still achieved 60.0%, demonstrating strong data efficiency (**Figure 2g**). These results highlight that FERAL achieves high and stable performance with minimal annotation effort.

The second benchmarking dataset, MABe22 (Multi-Agent Behavior 2022) [42], is a large-scale, multi-species collection of datasets, each annotated across diverse behavioral categories. Among the MABe22 datasets, we evaluated FERAL on the rove beetle (*Sceptobius lativentris*) subset, comprising a beetle interacting one-on-one with its host ant (*Liometopum occidentale*). We focused on this subset because it pairs raw video with fully released, frame-level multiclass behavior annotations for a non-mammalian, symbiotic interaction. For the rove-beetle subset of MABe22, we evaluated FERAL’s frame-level predictions against expert annotations using the macro-averaged (macro) F1 score, following the evaluation protocol of the original publication. Macro-averaged F1 score is defined as the average of F1 scores for each behavior, where F1 is

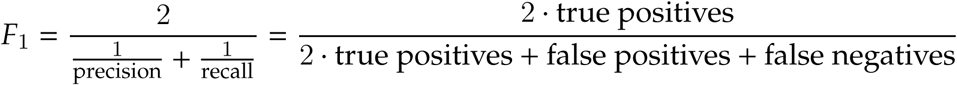

On MABe22, a random classifier achieves a macro-averaged F1 score of 0.24, while a perfect classifier achieves a macro-averaged F1 score of 1.0. The top-performing submission in the MABe22 challenge (Multimodal MoCo / SimCLR) achieved a macro F1 of 0.758 (Sun et al. 2023, Table 5) [42], which we adopt as the reference. Note that in the competition, the participants did not have access to the labeled data for the target behaviors during model training, and the labels were predicted by training a linear probe on top of frozen embeddings. So FERAL, which is trained directly on the target behavior labels, is expected to achieve higher performance. FERAL achieved macro F1 of 0.92, extending high performance from mammalian social behavior to multi-animal interactions between rove beetles and ants. Although this comparison is therefore not directly equivalent to the original MABe22 challenge protocol, it demonstrates that FERAL achieves high performance when trained directly on the available target-behavior labels.

We further benchmarked FERAL against DeepEthogram[24], the closest existing tool for direct video-to-behavior classification. Using DeepEthogram’s datasets and its headline metric, the frame-level F1 score (micro-averaged over all frames; for FERAL’s single-label predictions this is equivalent to top-1 accuracy), we reconverted six of its datasets (Mouse-Social, Mouse-Homecage, Mouse-Openfield, Mouse-Ventral1, Mouse-Ventral2, and Fly) into FERAL’s input format and trained a FERAL model on each of them. The DeepEthogram frame-level F1 values used in this comparison were read from Bohnslav et al. 2021 (Figure 3) [24] and cross-checked against the per-dataset scores kindly provided by the original authors.

**Figure 3:**
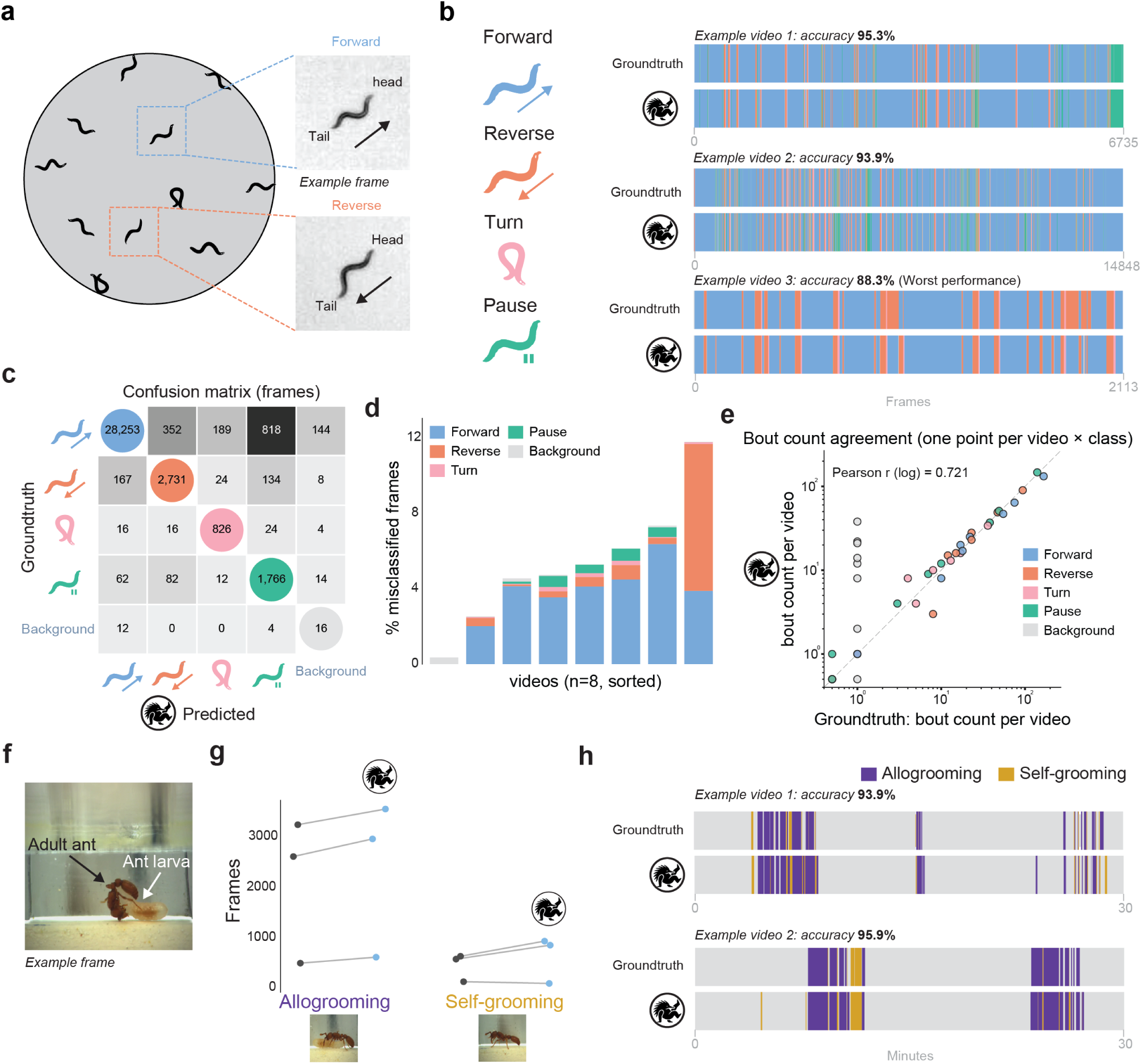
FERAL captures the temporal and visual appearance of behaviors across species and recording modalities. **(a)** Example frames from the *C. elegans* dataset showing forward and reverse locomotion states. **(b)** Representative frame-level behavioral time-series comparing ground-truth annotations (top) and FERAL predictions (bottom) for *C. elegans* locomotory states. **(c)** Confusion matrix of frame-level predictions versus heuristic-derived ground truth across the four locomotor states (*forward*, *reverse*, *turn*, *pause*). **(d)** Per-class fraction of incorrectly classified frames per video for *C. elegans*. **(e)** Agreement between ground-truth and predicted per-behavior bout counts across *C. elegans* videos (one point per video and class). **(f)** Example frame from recordings of pair interactions between an adult and larval clonal raider ant (*Ooceraea biroi*). **(g)** Quantitative comparison of ground-truth and predicted frame counts for each grooming category. **(h)** Representative frame-level behavioral time-series from *O. biroi* videos comparing ground-truth and FERAL predictions for self-grooming and allogrooming events.

FERAL matched or exceeded DeepEthogram’s frame-level F1 on five of six datasets, trailing only on Mouse-Social by 0.024 (**Table 1**). Bohnslav et al. 2021 [24] report a per-frame human-versus-consensus agreement for their five mouse datasets; FERAL exceeded it on four (Mouse-Homecage, Mouse-Openfield, Mouse-Ventral1, and Mouse-Ventral2), and on Mouse-Social its score (0.896) fell just short of both DeepEthogram and the human ceiling (∼0.92).

**Table 1:** FERAL versus DeepEthogram on the six original DeepEthogram datasets. Values are frame-level F1 (micro-averaged over all frames); for FERAL’s single-label predictions this equals top-1 accuracy. Human-vs-consensus is the per-frame human agreement reported by Bohnslav et al. 2021 (available for the five mouse datasets only). Bold marks the higher value in each pairwise FERAL–DeepEthogram comparison. FERAL values were regenerated from the reconverted datasets using stratified splits in which every behavior is represented in the training and test partitions; Mouse-Ventral1 reuses the original DeepEthogram split, and Mouse-Ventral2 and Mouse-Social report the mean of three stratified splits (see Methods).

| Dataset | FERAL | DeepEthogram | Human-vs-consensus |
| --- | --- | --- | --- |
| Mouse-Social | 0.896 | **0.920** | $\geq 0.910$ |
| Mouse-Homecage | **0.905** | 0.760 | $\geq 0.700$ |
| Mouse-Openfield | **0.883** | 0.870 | $\geq 0.790$ |
| Mouse-Ventral1 | **0.943** | 0.910 | $\geq 0.900$ |
| Mouse-Ventral2 | **0.978** | 0.970 | $\geq 0.960$ |
| Fly | **0.950** | 0.940 | — |

To assess whether FERAL’s residual errors track the disagreement between annotators rather than systematic bias, we stratified Mouse-Ventral1 test frames by per-frame inter-annotator agreement and evaluated a FERAL model trained on annotator 1’s labels against those same annotator 1 labels ([24]). Accuracy with respect to annotator 1’s labels fell as human agreement fell: 98.9% on frames where all three annotators agree, 94.0% where two of three agree, and only 27.5% on the 0.7% of frames where all three disagree (**Figure 2h**). Because accuracy is scored against annotator 1’s labels, the low value on the fully-disagreed frames is expected—these are the most ambiguous frames, where even annotator 1’s own label is unreliable—and shows that model error tracks genuine human label noise rather than introducing systematic annotator-specific bias. A complementary cross-annotator analysis on the same Mouse-Ventral1 data showed that model–annotator agreement tracked, and stayed at or just below, human–human agreement across every annotator pair: for the two concordant annotators (1 and 2), a FERAL model trained on annotator 1 reached a macro F1 of 0.59 against annotator 2, close to the 0.61 the two humans achieved with each other, whereas agreement with the more idiosyncratic annotator 3 was similarly low for both the model (0.32–0.37) and the humans (0.38–0.43) (**Extended Figure 1d**). FERAL therefore learns the cross-annotator consensus rather than one annotator’s idiosyncrasies.

To translate model performance into annotator effort, we reframed the data-efficiency analysis at the level of annotated behavioral bouts. On the two most heterogeneous DeepEthogram datasets, Mouse-Social and Mouse-Ventral1, we randomly retained 75%, 50%, 25%, and 12.5% of the annotated bouts, removing the rest from training, and retrained FERAL with otherwise identical hyperparameters. FERAL retained near-full macro F1 with 75% of the annotated bouts (within ∼5% of the full-data F1 on both datasets), and 90–95% at 50%. Below 25%, the rarest classes collapsed to zero positives predicted (**Figure 2i**). In practical terms, annotation budgets can be halved on common behaviors without measurable cost, while rare classes require more extensive annotation to avoid the collapse observed at low sampling rates.

Finally, we probed how FERAL behaves on behaviors absent from its training schema by training four leave-one-out variants on Mouse-Ventral1, each removing one foreground class (face-grooming, body-grooming, digging, scratching) and relabeling its frames as background. Across all four variants, held-out frames migrated either to a semantically related foreground class (e.g., face-grooming → body-grooming, 55%; scratching → body-grooming, 41%) or to background (digging and body-grooming migrated almost entirely to background, 100% and 96%), never to a distant unrelated class; retained-class F1 stayed within 0.11 of the full-schema baseline and background F1 remained ≥ 0.964 (**Extended Figure 1c**). Being a supervised method, FERAL does not discover novel behaviors, and behavioral discovery requires unsupervised pipelines such as MoSeq [43], keypoint-MoSeq [19], B-SOiD [20], or VAME [21]).

### FERAL captures the temporal structure and visual appearance of behaviors

To assess FERAL’s ability to generalize across species and recording modalities, we applied it to two datasets, both acquired in laboratory conditions, representing distinct levels of behavioral organization and scene complexity.

#### Single-animal behavior in *C. elegans* nematodes

This dataset comprises recordings of freely moving *C. elegans* nematodes performing four canonical locomotor behaviors: forward crawling, reverse crawling, turning, and pausing (**Figure 3a**) [44–48]. Unlike other datasets used in this study, these behavioral labels were not manually annotated; instead, each frame was automatically assigned to a locomotory state using a rule-based heuristic pipeline adapted from prior work [48, 49]. In this pipeline, worms were segmented from the background, centroid positions were extracted, head–tail orientation was inferred from midline dynamics, and locomotor state was determined using speed thresholds, direction of motion, and self-intersection criteria. This is therefore a benchmark of model agreement with an algorithmic gold standard rather than with manual expert annotation. The resulting behavioral classification was verified by an expert annotator on a subset of frames (**Extended Figure 2b**).

For each original video with many worms, the centroid of each worm was continuously tracked, and a cropped image window centered on the animal was extracted to maintain consistent framing. FERAL leverages temporal context by integrating information from preceding and subsequent frames to classify each moment in time. As a result, it accurately segmented all four behavioral states and correctly distinguished between forward and reverse locomotion. These two locomotory states appear nearly identical in single frames but become separable through their temporal dynamics (**Figure 3b, Supplementary Video 2**).

Confusion-matrix analysis (**Figure 3c**) showed that the majority of errors were confined to transitions between kinematically adjacent states, such as “forward” versus “pause” or “forward” versus “reverse.” Across videos, FERAL maintained high frame-level accuracy (overall ∼94% against the algorithmic ground truth; **Figure 3d**), and predicted per-behavior bout counts closely tracked the algorithmic ground truth (**Figure 3e**; per-video accuracy and bout-duration distributions in **Extended Figure 2a,c**). Beyond its strong performance, this analysis illustrates how FERAL can be seamlessly integrated into existing behavioral pipelines as a post-tracking module, complementing traditional centroid-based approaches by directly mapping cropped video segments to behaviors.

#### Pair interactions in *Ooceraea biroi* clonal raider ant

We next tested FERAL on a dataset capturing *O. biroi* adult-larva interactions, which included annotations (J.Z.) for self-grooming and allogrooming events (**Figure 3f**). Self-grooming involves individuals cleaning their own body, while allogrooming targets another colony member which, in this case, is a larva. Accurate classification thus requires recognizing not only motion patterns but also the spatial relationship between the adult and the larva. This poses a major challenge for pose-based pipelines: (i) pose estimation is unreliable under frequent occlusions; and (ii) modeling these social interactions using skeletonized data requires integrating information from multiple individuals [50–52].

FERAL bypasses the need for pose estimation and pose-based segmentation, thus avoiding these challenges altogether. It reliably identified both self- and allogrooming events directly from raw videos (**Supplementary Video 3**), maintaining the overall temporal dynamics of each behavior across videos with high fidelity to the original expert annotations (**Figure 3f-h**; confusion matrix in **Extended Figure 2d**).

Altogether, these results highlight FERAL’s ability to extract meaningful behaviors from spatiotemporal patterns that are difficult to access with single-frame image models or pose-based approaches, demonstrating its capacity to generalize from individual locomotion to social behaviors.

### FERAL generalizes to field recordings of wild animals

Given FERAL’s robust performance on datasets acquired in the laboratory, we next evaluated its generalization to field recordings, which typically exhibit higher scene complexity than laboratory videos.

#### Vigilance behavior in zebras

This dataset consisted of aerial videos of wild Grevy’s zebras (*Equus grevyi*) recorded at the Mpala Research Center in Laikipia, Kenya using a drone equipped with a camera. Free-ranging groups of zebras were filmed from directly overhead. As with the *C. elegans* dataset, the centroid of each animal was continuously tracked following methods described in [54] and a cropped image window was centered on the animal to maintain consistent framing. Individual videos were annotated in BORIS by an expert (B.R.C.) to identify bouts of vigilance behavior, defined as the individual standing still with its head raised (**Figure 4a**). Due to the overhead perspective, the vertical head position of the zebras is challenging to detect. We initially attempted to identify vigilance bouts using pose estimation: nine keypoints (nose, head, base of neck, left and right shoulders, left and right hips, and base and tip of tail) were annotated in 900 cropped images and used to train a DeepPoseKit [18] pose-estimation model, which was then applied to a subset of cropped videos of individual zebras. These pose estimates did not support reliable vigilance detection. We confirmed this quantitatively: gradient-boosted classifiers (XGBoost [55], CatBoost [56]) trained on the DeepPoseKit keypoints collapsed to near-chance performance when required to generalize to a held-out recording (balanced accuracy ≈ 0.58, chance = 0.50; macro-F1 ≈ 0.55), well below FERAL’s macro-F1 of 0.785 on the same task from raw video (**Extended Figure 3a**).

**Figure 4:**
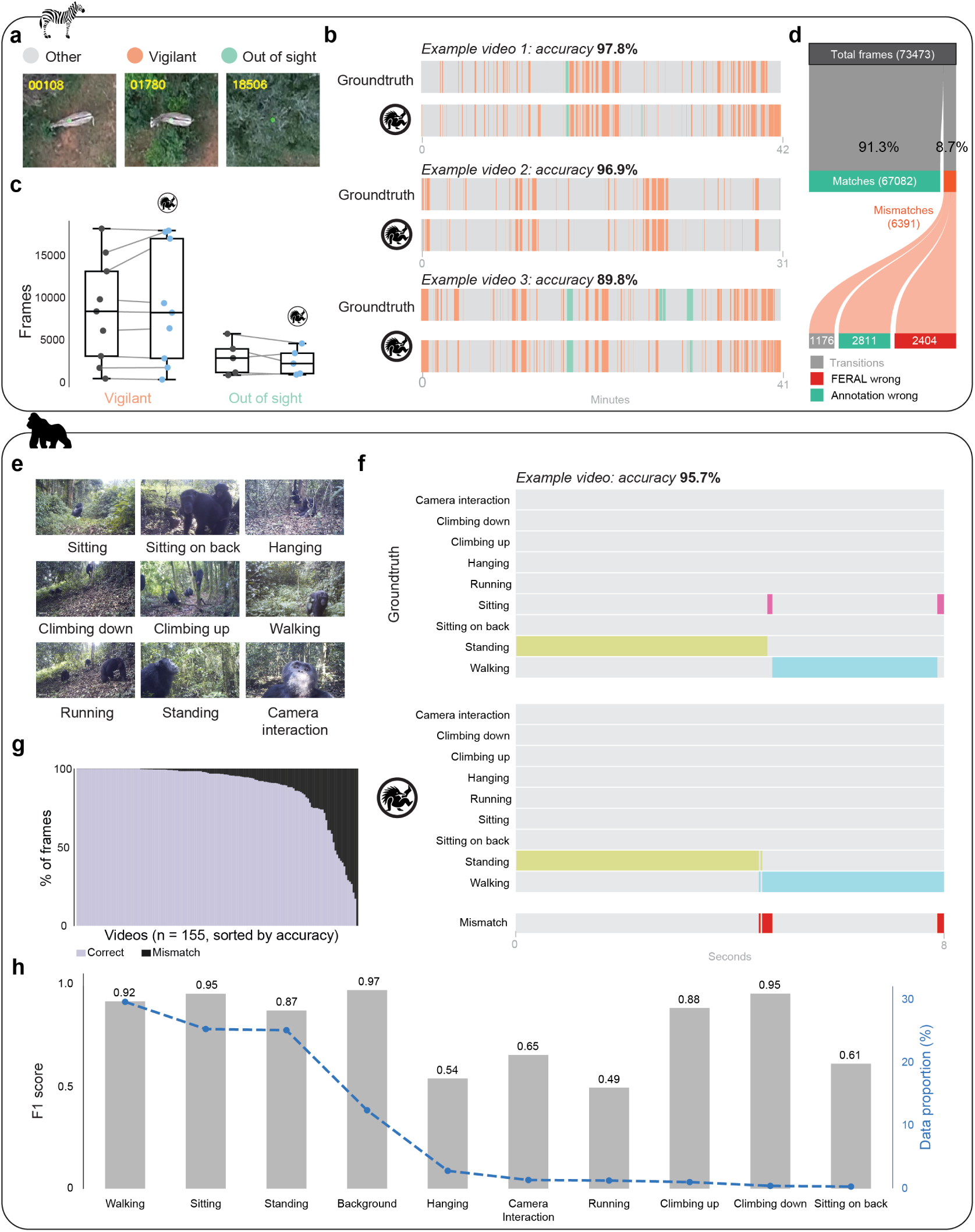
FERAL generalizes to field recordings of wild animals. **(a)** Representative drone frame of wild Grevy’s zebras (*Equus grevyi*), Mpala Research Center, Kenya. **(b)** Example frame-level behavioral sequences (expert annotation, top; FERAL, bottom) for three zebra videos. **(c)** Per-video number of frames assigned to the *vigilant* and *out-of-sight* states, expert annotation versus FERAL. **(d)** Frame-by-frame manual audit (B.R.C.) of all FERAL–annotator mismatches in one held-out zebra video. **(e)** Example PanAf500 [53] frames (wild *Pan troglodytes* and *Gorilla gorilla*, camera traps), one per annotated class. **(f)** Example frame-level ethogram (expert, top; FERAL, bottom) for PanAf500. **(g)** Per-video fraction of correctly (light) and incorrectly (dark) classified frames, PanAf500 (n = 155 videos). **(h)** Per-class F1 (bars) and labeled-data proportion (line) for PanAf500; classes ordered by descending data proportion.

FERAL accurately detected the onset and duration of vigilance periods directly from raw video, closely matching expert annotations across most recordings. The fraction of correctly labeled frames was high across recordings, at or above 88% in eight of nine videos, with the shortest recording reaching 67% (**Figure 4b**, **Extended Figure 3b**). Comparison of the total number of frames assigned to the *vigilant* and *out-of-sight* states per video showed strong agreement between FERAL and expert annotation (**Figure 4c**), indicating that FERAL generalized to aerial perspectives under the natural variability of field conditions; the corresponding per-state confusion matrix (*vigilant* recall 87%, *out-of-sight* recall 67%) is shown in **Extended Figure 3c**.

To characterize the nature of FERAL’s residual errors, an expert annotator (B.R.C.) manually reviewed every frame-level disagreement between FERAL and the manual annotation in a representative held-out test video (73,473 frames; **Figure 4d**). FERAL’s prediction matched the annotation on 67,082 frames (91.3%); of the 6,391 mismatched frames (8.7%), only 2,404 (37.6%) reflected genuine misclassifications by FERAL. The remainder arose from label noise rather than model error: 1,176 frames fell within behavioral transitions whose exact timing was not annotated frame-precisely, and 2,811 frames were cases in which FERAL was correct and the manual annotation was in error. Among FERAL’s genuine errors, the dominant confusions were *other* scored as *vigilant* (1,163 frames) and *vigilant* scored as *other* (892 frames). Accounting for annotation errors and transition-timing differences raises the effective frame-level accuracy on this video from 91.3% to approximately 96.7% (**Figure 4d**).

#### Chimpanzee and gorilla behavior in the wild

To further assess FERAL’s generalization to complex, naturalistic scenes, we evaluated its performance on the PanAf500 (Pan African Programme 500-clip subset) dataset [53], which contains camera-trap videos of wild chimpanzees (*Pan troglodytes*) and gorillas (*Gorilla gorilla*) recorded across multiple African field sites as part of the Pan African Programme: The Cultured Chimpanzee [57]. In each clip, all individuals were detected with a bounding box and each individual’s behavior was manually annotated with fine-grained behavioral categories, including locomotor (e.g., walking, climbing, running), postural (e.g., standing, sitting, hanging), and interaction-related behaviors (**Figure 4e**), testing FERAL’s capacity to handle visual and behavioral complexity on a multi-species dataset.

Camera-trap footage varied substantially in lighting, vegetation, and visibility. FERAL reached top-1 accuracy of 90.8% and macro averaged recall of 81.0% over the nine classes with positives in the official PanAf500 test split following the original testing protocol of [53] (**Figure 4f-h, Supplementary Video 4**, **Extended Figure 3d**). FERAL exceeded all the methods reported by Brookes et al. 2024 [53] (X3D, I3D, 3D ResNet-50) on both metrics. It also surpassed recent primate-specific methods (ChimpVLM [26], DAP [25], PriVi [27]) (**Table 2**). Among the nine test-positive classes, performance was highest on prevalent behaviors including *walking*, *sitting* and *standing* and was lowest on rare ones: *camera interaction*, *running* (**Figure 4h**).

**Table 2:** PanAf500 behavior-recognition comparison. Top-1 accuracy and class-averaged recall over the test-positive classes of the official PanAf500 test split. X3D, I3D, and 3D ResNet-50 are the 3D-CNN baselines of Brookes et al. 2024 [53].

| Method | Top-1 acc. (%) | Class-avg recall (%) |
| --- | --- | --- |
| X3D [53] | 80.0 | 50.4 |
| I3D [53] | 79.3 | 42.2 |
| 3D ResNet-50 [53] | 77.5 | 55.2 |
| ChimpVLM [26] | 84.9 | 61.9 |
| DAP [25] | 87.2 | 56.4 |
| PriVi [27] | 86.7 | 62.8 |
| **FERAL** | **90.8** | **81.0** |

Together, these results demonstrate that FERAL’s performance on videos acquired in laboratory settings transfers robustly to field recordings spanning multiple species (zebras, chimpanzees, and gorillas), habitats, and acquisition modalities. Its ability to extract meaningful behavioral structure directly from raw video highlights the model’s versatility and scalability for quantifying natural behavior in ecologically realistic contexts.

### FERAL identifies emergent collective behaviors directly from raw video

To test whether FERAL generalizes from individual and pair interactions to collective dynamics, we applied it to recordings of clonal raider ant (*Ooceraea biroi*) colonies filmed continuously over several days (**Figure 5a**). In this species, foraging occurs in collective group raids [58]. Manual annotation of such raiding events was used as ground truth. Despite the high density of individuals, FERAL accurately detected the onset and duration of collective raids directly from raw video frames (**Figure 5b**). FERAL captured these events without requiring individual tracking or explicit modeling of group structure. By contrast, conventional multi-animal tracking approaches would need to reconstruct trajectories for each individual [17, 59] and subsequently infer collective states through post-tracking analyses [58]. FERAL bypasses these steps, enabling scalable, tracking-free quantification of collective behavior.

**Figure 5:**
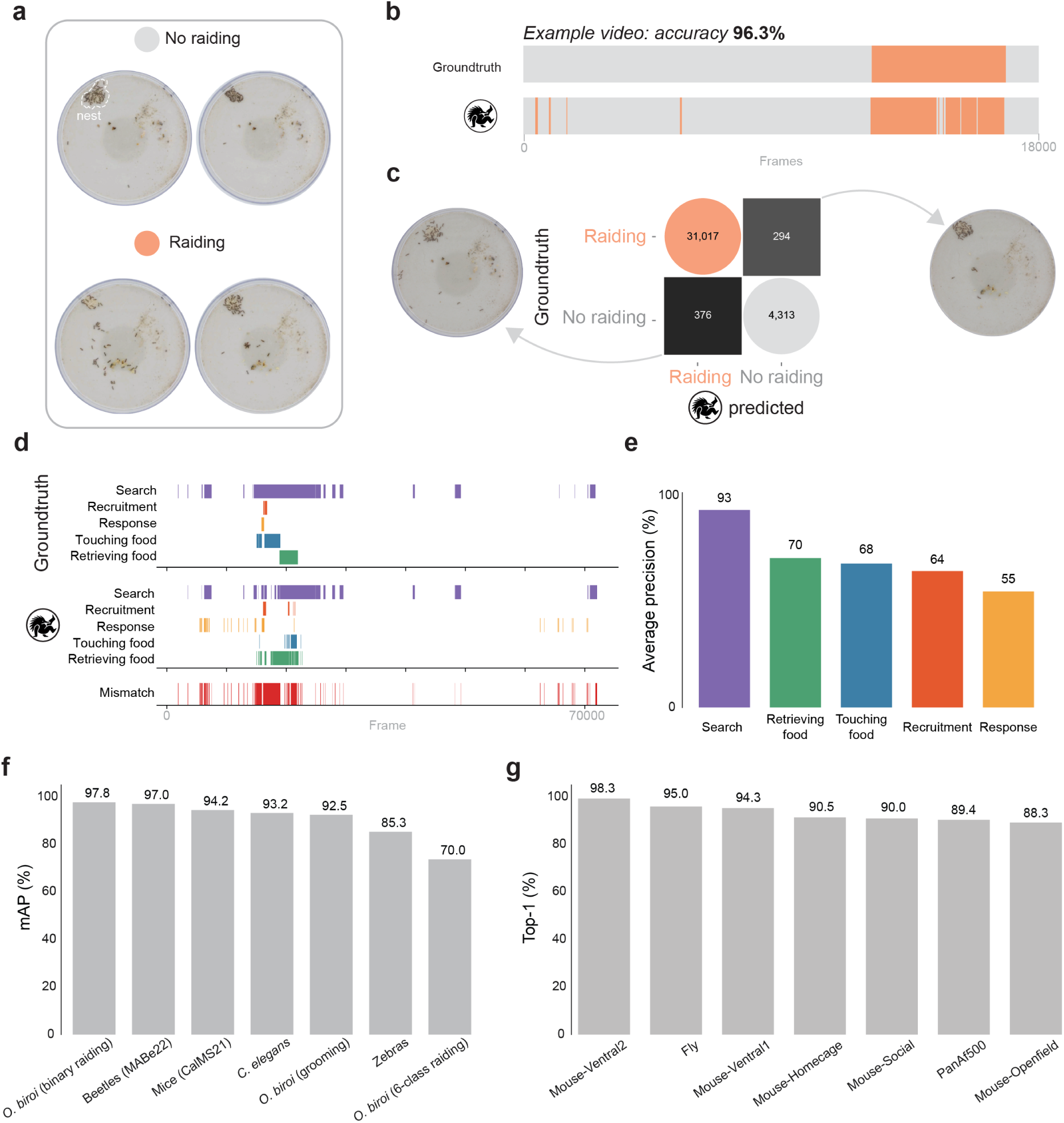
FERAL captures emergent collective behavior and achieves high performance across datasets. **(a)** Representative frames from recordings of clonal raider ant (*Ooceraea biroi*) colonies during non-raiding (top) and raiding (bottom) phases, with nests indicated by dashed outlines. **(b)** Example frame-level time-series of *binary* raid detection (raiding vs. non-raiding) in the colony dataset, comparing ground-truth annotation (top) and FERAL prediction (bottom). **(c)** Confusion matrix for binary raid detection across all colony videos; insets show representative frames illustrating the visual appearance of misclassified frames. **(d)** Example frame-level time-series of the *five-class* raid substructure in a separate, spatially structured arena dataset, comparing ground-truth annotation (top) and FERAL prediction (bottom); in contrast to the binary detection in (b), this resolves individual ant behaviors and collective raid phases within a raid. **(e)** Per-state average precision for the five raid-substructure classes. **(f)** mAP of FERAL across all reported datasets. **(g)** Top-1 performance of FERAL across reported datasets.

Confusion-matrix analysis (**Figure 5c**) revealed that misclassified “raiding” frames were not random but systematically associated with moments of intense movement, when many ants had exited the nest and were exploring the arena. Conversely, most “no raiding” misclassifications occurred during phases of a raid when most ants were in the nest and overall activity declined. These patterns suggest that FERAL’s classifications are driven by the colony-level motion and spatial distribution of individuals, demonstrating that the model learns the collective visual signature of raiding behavior directly from scene appearance.

By recognizing colony-level behaviors directly from video, FERAL extends the reach of direct video-to-behavior analysis to emergent social dynamics that arise from distributed coordination among many animals. This ability opens new possibilities for studying the neural, ecological, and evolutionary principles governing collective behavior across species and environments.

Detecting whether a raid is underway is a coarse, binary description of collective state. We next asked whether FERAL could resolve the finer internal organization of a raid. For this we analyzed a separate dataset of *O. biroi* raids recorded in larger, spatially structured arenas (small colonies foraging from a nest into a connected arena; see Methods), in which we could annotate both individual ant behaviors (*touching food*, *retrieving food*) and collective raid phases (*search*, *recruitment*, *response*), and trained FERAL to predict these five behavioral states directly from raw video.

FERAL recovered the temporal structure of the raid, with predicted state sequences broadly tracking the annotated behavioral sequences (**Figure 5d**). Across the five states, FERAL reached a mAP of 70.0% (**Figure 5f**), with the highest per-state average precision for *search* (0.93) and the lowest for *response* (0.55) (**Figure 5e**).

Because raids vary considerably in duration and clarity across recordings, multiclass state recognition is markedly harder than binary raid detection (macro-averaged mAP 70.0% versus 97.8% for binary raiding). We therefore present it as a proof of concept that direct video-to-behavior analysis can begin to resolve the substructure of collective dynamics (**Figure 5f**).

### FERAL runs efficiently across hardware configurations

FERAL is designed to run on a broad range of hardware, so that compute is not a barrier to adoption. We benchmarked training and inference throughput for the default and lite configurations across 8 single-GPU setups, spanning data-center accelerators (H100, A100, L40S) and consumer cards (RTX 5090, RTX 3090) (**Table 3**). FERAL also provides a gradient checkpointing option (gc), which slows down training by around 20% but reduces the VRAM footprint by around 3x.

**Table 3:** FERAL training and inference cost across GPU configurations. GPU costs are provided for the datacenter and community options.

| GPU | Config | VRAM<br>(GB) | \$/h<br>(dc.) | \$/h<br>(com.) | Train<br>(ch/s) | Inference<br>(ch/s) | Full CalMS21<br>(10 ep) | Inference 1 h<br>30-fps video |
| --- | --- | --- | --- | --- | --- | --- | --- | --- |
| H100 SXM | default | 80 | \$3.29 | \$2.00 | 14.9 | 24.6 | 3.0 h (\$9.8/\$6.0) | 2.3 min |
| RTX 5090 | default | 32 | \$0.99 | \$0.40 | 6.3 | 12.2 | 7.0 h (\$6.9/\$2.8) | 4.6 min |
| A100 SXM | default | 80 | \$1.49 | \$0.60 | 6.2 | 11.6 | 7.2 h (\$10.7/\$4.3) | 4.8 min |
| L40S | default | 48 | \$0.86 | \$0.80 | 6.2 | 11.5 | 7.1 h (\$6.1/\$5.7) | 4.9 min |
| RTX 3090 | default | 24 | \$0.46 | \$0.14 | 2.4 | 4.8 | 18.3 h (\$8.4/\$2.6) | 11.8 min |
| H100 SXM | lite | 80 | \$3.29 | \$2.00 | 20.3 | 38.6 | 2.2 h (\$7.2/\$4.4) | 1.5 min |
| RTX 5090 | lite | 32 | \$0.99 | \$0.40 | 10.1 | 25.2 | 4.4 h (\$4.3/\$1.8) | 2.2 min |
| A100 SXM | lite | 80 | \$1.49 | \$0.60 | 9.5 | 21.0 | 4.7 h (\$6.9/\$2.8) | 2.7 min |
| L40S | lite | 48 | \$0.86 | \$0.80 | 9.7 | 24.4 | 4.6 h (\$3.9/\$3.7) | 2.3 min |
| RTX 3090 | lite | 24 | \$0.46 | \$0.14 | 4.2 | 12.3 | 10.6 h (\$4.9/\$1.5) | 4.6 min |
| H100 SXM | default+gc | 80 | \$3.29 | \$2.00 | 12.9 | 21.6 | 3.4 h (\$11.3/\$6.9) | 2.6 min |
| RTX 5090 | default+gc | 32 | \$0.99 | \$0.40 | 5.5 | 11.6 | 8.0 h (\$7.9/\$3.2) | 4.9 min |
| A100 SXM | default+gc | 80 | \$1.49 | \$0.60 | 5.3 | 11.2 | 8.4 h (\$12.5/\$5.0) | 5.0 min |
| L40S | default+gc | 48 | \$0.86 | \$0.80 | 5.1 | 11.3 | 8.7 h (\$7.5/\$6.9) | 5.0 min |
| RTX 3090 | default+gc | 24 | \$0.46 | \$0.14 | 2.0 | 4.8 | 21.8 h (\$10.0/\$3.0) | 11.8 min |
| RTX 3080 | default+gc | 10 | — | \$0.36 | 1.85 | 4.5 | 23.9 h (\$8.61) | 12.6 min |
| RTX 3070 | default+gc | 8 | — | \$0.11 | 1.25 | 3.0 | 35.4 h (\$3.89) | 18.5 min |
| RTX 3060 | default+gc | 12 | — | \$0.07 | 0.76 | 1.8 | 57.9 h (\$4.06) | 30.7 min |
| RTX 3080 | lite+gc | 10 | — | \$0.36 | 2.67 | 10.6 | 16.5 h (\$5.94) | 5.3 min |
| RTX 3070 | lite+gc | 8 | — | \$0.11 | 1.82 | 7.0 | 24.2 h (\$2.66) | 8.0 min |
| RTX 3060 | lite+gc | 12 | — | \$0.07 | 1.15 | 4.5 | 38.4 h (\$2.69) | 12.5 min |

For each GPU and configuration, we measured throughput by running the complete FERAL training pipeline—identical to a real run, including mixup, gradient-norm logging, the AdamW optimizer with cosine schedule, and the compiled forward/backward pass. We ran five epochs over a fixed synthetic dataset of 100 training batches (batch size 4) and 100 validation batches (batch size 8), reporting the throughput of the final, steady-state epoch and discarding the earlier epochs, which absorb compilation and dataloader warm-up. GPU clock and power draw were sampled throughout and per-epoch wall-clock was stable across the steady-state epochs (no upward drift indicative of thermal throttling), with cards held near their sustained power limits. All benchmarks used PyTorch 2.11.0 with the CUDA 12.8 (cu128) build, installed identically on every GPU. Host driver versions varied but the PyTorch CUDA runtime was uniform across cards. We noticed that using the latest PyTorch version sped up training and inference. Training throughput is taken from the training-epoch wall-clock and inference throughput from the validation-epoch wall-clock. Because video decoding is negligible at 256 by 256 resolution, GPUs remained at 100% utilization throughout. We estimate full ten-epoch training run times on CalMS21 by dividing 15,768 training chunks per epoch × 10 epochs by the estimated training speed, and adding 4,084 test chunks divided by the estimated inference speed. Despite excluding some overheads, these estimates match the actual runtimes of our experiments on H100 and L40S GPUs.

To estimate GPU costs, we report two price tiers: datacenter and community. For datacenter pricing, we use *RunPod Secure Cloud* prices. For community pricing, we use *VAST.ai*, a marketplace where independent hosts rent out their private GPUs. All prices are reported as of June 19, 2026.

For *VAST.ai*, we report the lowest price we could find among hosts that met our minimum requirements: reliability ≥ 0.97, at least 16 effective CPU cores, 32 GB RAM, 50 GB disk space, and 200+ Mbps download speed. These filters remove unreliable hosts and ensure enough CPU, memory, storage, and network bandwidth for training without obvious bottlenecks.

This lets us take advantage of the larger supply of consumer GPUs. While median community prices are often close to datacenter prices, the open-market structure means that the lowest available prices can be much lower, especially for popular GPUs with abundant supply like an RTX 5090.

We chose these GPUs because FERAL currently requires an NVIDIA GPU from the Ampere generation or newer (compute capability ≥ 8.0, required for BF16 and FlashAttention). Older accelerators available for free on Google Colab, such as the T4 and V100, are not supported by FERAL. FERAL can therefore be run locally on a single workstation GPU, on an institutional high-performance cluster, or on cloud GPU platforms such as RunPod or Google Colab Pro for users without access to local resources. For these users, we provide step-by-step deployment guides and a ready-to-run notebook.

A full default CalMS21 run takes under 4 hours on an H100 and around 7 hours on midrange cards (A100, L40S, RTX 5090), with costs around $6-11 per run for datacenter GPUs and $2-6 for community GPUs. The most cost-efficient GPU turned out to be the RTX 5090, a latest-generation prosumer GPU, enabling a full *lite* run on a vast.ai machine for $1.50. Inference runs faster than real time on every GPU tested: one hour of 30-frames-per-second video is processed in ∼2.3 minutes on an H100 and ∼30 minutes on an RTX 3060.

The *lite* option provides a faster and more affordable option for experiments on smaller GPUs, although the default option can also run on smaller GPUs with gradient checkpointing. While full training may not be fully practical on those smaller GPUs, it can still work if users run fewer training epochs or have access to multiple smaller GPUs.

## DISCUSSION

The quantitative study of animal behavior has advanced rapidly with the integration of computer vision and deep learning [3, 7]. Existing methods have excelled at answering where animals and their body parts are over time, yet determining what actions animals are performing remains challenging, especially in complex environments. This limitation constrains discovery across the fields of ethology, neuroscience, and ecology [8, 12], where understanding behavior and its structure is essential.

FERAL addresses this gap by reframing behavioral quantification as a video-understanding problem. Instead of relying on intermediate abstractions such as pose or trajectory, FERAL maps raw video directly to frame-level behavioral labels, capturing both the temporal and visual appearance of animal actions. It achieves high performance across datasets, detecting discrete actions with high precision on a wide range of scene complexities (**Figure 5f-g**). This generalization is enabled by its design. Its foundation-model backbone allows robust fine-tuning even with limited behavioral annotations [32].

FERAL sits within a broader landscape of automated behavioral quantification. The closest pixel-input precedent is DeepEthogram [24], which performs supervised classification of behavior directly from raw video using a two-stream convolutional neural network (CNN). FERAL extends this approach with a fine-tuned video foundation model and substantially improves accuracy on five of six DeepEthogram datasets. For unsupervised behavioral discovery — uncovering categories not specified by a human annotator — methods such as MoSeq [43], keypoint-MoSeq [19], B-SOiD [20], and VAME [21] remain the appropriate tools. FERAL is complementary to, not a substitute for, this line of work. Recent video-foundation-model approaches for animal behavior — DAP [25], ChimpVLM [26], and PriVi [27] — target behaviors only for primates. FERAL is a general-purpose drop-in supervised pipeline that runs across species with frame-level labels alone.

FERAL operates on user-supplied behaviors and emits per-frame predictions constrained by this list of behaviors. It does not autonomously discover new behavioral categories.

Despite these limitations, FERAL is designed for broad accessibility: it is an open-source, freely available toolkit (getferal.ai) that includes comprehensive documentation, tutorials, and benchmark datasets, lowering the technical barrier for users across disciplines. Its modular architecture supports all stages of analysis, from preprocessing and training to inference, and can be deployed locally on desktop computers, on high-performance clusters, or through cloud-based GPU services such as Google Colab Pro and RunPod.

FERAL integrates with existing pipelines. By default, FERAL focuses on frame-level segmentation and does not assign persistent identities. Nevertheless, it can be paired with multi-animal tracking systems such as TRex [17], which produce animal-centered video segments suitable for direct inference. Together, these tools provide a unified solution to the two central questions of behavioral quantification: where animals are and what actions they are performing.

FERAL is a supervised classifier: it can only predict behaviors present in its training labels, and frames whose behavior is absent from the schema are routed to the nearest semantic neighbor or to background rather than discovered as new; behavioral discovery remains the domain of unsupervised methods such as MoSeq [43], keypoint-MoSeq [19], and B-SOiD [20]. Additionally, FERAL outputs categorical labels, not continuous kinematic variables (joint angles, gait phase, head orientation); pose-tracking pipelines such as DeepLabCut [14], SLEAP [15], and LightningPose [16] remain the standard approach for postural readouts, and the cleanest analyses will often combine them with FERAL (pose for kinematics, FERAL for behavioral bouts).

Importantly, FERAL classifies the scene rather than the individuals in it: in pair and social settings the model does not assign actor–recipient identity. In scenarios in which the animal of interest occupies only a small fraction of the frame (single-worm assays, single-zebra vigilance, multi-ape camera-trap clips), FERAL requires an upstream tracker or detector to produce animal-centered crops. This requirement is shared with most supervised pose-based and pixel-based classifiers.

FERAL shifts annotation effort to the training stage rather than eliminating it: a moderate annotation budget (∼50% of bouts) recovers most of the benefit for common behaviors, but rare-class detection remains bound by the labeling effort directed at that specific class.

By combining strong performance with user-friendly design, FERAL establishes a robust foundation for behavioral segmentation across species and experimental paradigms. FERAL code and documentation are available at `getferal.ai` (Zenodo DOI to be assigned).

Since this work was released as a preprint, FERAL has been adopted by many research groups on more than 250 datasets across multiple species and experimental paradigms, and we are hopeful that this tool will be helpful to the scientific community.

## METHODS

### Datasets

We evaluated FERAL across 14 datasets spanning a range of species, behavioral contexts, and recording modalities. All datasets provided raw video data paired with frame-level behavioral annotations, enabling direct video-to-behavior mapping. CalMS21, MABe22, PanAf500 and DeepEthogram datasets were obtained from public sources. We did not perform new vertebrate experiments on these data, and the original ethics approvals are reported in the source publications. Original ethics approvals for our new datasets are stated in the corresponding subsection. Other open-source datasets containing only keypoint trajectories or bounding boxes without frame-level behavioral labels were excluded, as the current implementation focuses on frame-level classification. Training and test dataset statistics, including the number of frames per split for each dataset, are reported in Table 10, and class distribution and per-class performance are given in Table 12. Performance metrics for each dataset are reported in Table 13. Note that for single-label classification, we include an “other” class that the model is trained to predict when no relevant behaviors are present. For all reported class-averaged metrics, the “other” class is excluded.

#### CalMS21 (Mouse social interactions)

The Caltech Mouse Social Interactions (CalMS21) dataset captures freely behaving mice (*Mus musculus*) engaged in resident–intruder assays [39]. Each recording includes frame-level labels for social behaviors such as attack, mount, and investigation. Following prior work, we evaluated performance using the mean average precision (mAP) metric (macro-averaged AP over foreground classes) on the same train–test split defined in the original challenge.

#### MABe22 (Multi-Agent Behavior 2022)

MABe22 [42] is a large-scale multi-species benchmark designed for evaluating multi-agent behavior analysis. It includes recordings of mouse triplets, rove beetles (*Sceptobius lativentris*), and *Drosophila melanogaster* flies paired with expert-annotated behavioral categories. As only the beetle subset provides raw video aligned with human frame-level labels, we restricted our evaluation to this dataset. Behavioral categories are *grooming object*, *grooming self*, *idle alone*, *idle object*, *exploring object*, *exploring alone* annotated by trained ethologists. We report macro-averaged F1 scores on the original test split from the paper.

The original MABe22 beetle partition is divided into four subsets: *user train* (released for unsupervised representation learning, without the six target-behavior labels), *evaluation train* (the labeled subset used to train the benchmark’s linear classifier), *test1*, and *test2*. However, in the publicly released data, only the *user train* can be distinguished from the rest. The boundaries between *evaluation train*, *test1*, and *test2* are not provided, so we could not reproduce the exact split used in the original benchmark. Furthermore, only 1122 of the 9491 evaluation clips (∼12%) carry per-frame annotations for behaviors. These labeled clips come from the combined *evaluation train*, *test1*, and *test2* splits. We therefore pooled all of them and divided them randomly into 80% training and 20% test.

Rather than training a linear classifier on top of frozen embeddings, as in the MABe22 competition, we fine-tune the model directly on these labeled clips.

#### PanAf500

PanAf500 [53] is a dataset of wild ape behavior, featuring both chimpanzees and gorillas recorded by camera traps across multiple African field sites within the Pan African Programme. The cameras operate during the day and at night, capturing a range of behaviors expressed by single individuals and groups under diverse weather and lighting conditions. The dataset documentation provides field collection and ethics context through the Pan African Programme [53, 57].

The original work introduces two datasets: PanAf20k, containing 20k clips with a single behavior label per clip, and PanAf500, consisting of 500 clips in which each individual ape is detected with a bounding box in every frame and assigned a behavior label. Because we wanted to evaluate FERAL on per-frame behavior classification, we used the smaller PanAf500 subset.

In our experiments, we retained the original held-out test split (75 15s videos) and constructed a single train set by combining the validation and train splits from the original paper (425 15s videos). We reformatted the per-frame bounding-box labels into per-individual-tracklet videos with one frame-level label per tracklet, in order to evaluate FERAL’s performance against a single target per frame. Three preprocessing choices slightly deviate from the official `single_ape.py` loader: (i) we use a square max(*w, h*) crop with 5% padding (the official loader uses an aspect-preserving crop), to match the V-JEPA 2 backbone’s square input shape; (ii) we gap-fill bounding boxes across short occlusion frames (up to 20 missing-detection frames per tracklet) and set the labels for occluded frames as background, so that the per-tracklet video is a contiguous sequence; (iii) we filter out tracklets visible in fewer than 16 video frames. The remaining tracklets are padded to a length that is a multiple of 64 frames using black frames and a background label. These deviations were required for the fixed input shape of the V-JEPA 2 backbone and can possibly slightly inflate FERAL’s reported accuracy relative to the Brookes et al. 2024 baselines (X3D, I3D, 3D ResNet-50), which use the official loader. The background class is then removed from test metric calculations.

Compared to the default FERAL setup that uses per-frame metrics, the original paper splits the bouts into chunks of 16 frames dropping the rest, and predicts a single label for such bouts. We report 2 sets of metrics per frame and per 16-frame chunk to be comparable with available benchmarks in Table 4.

**Table 4:**
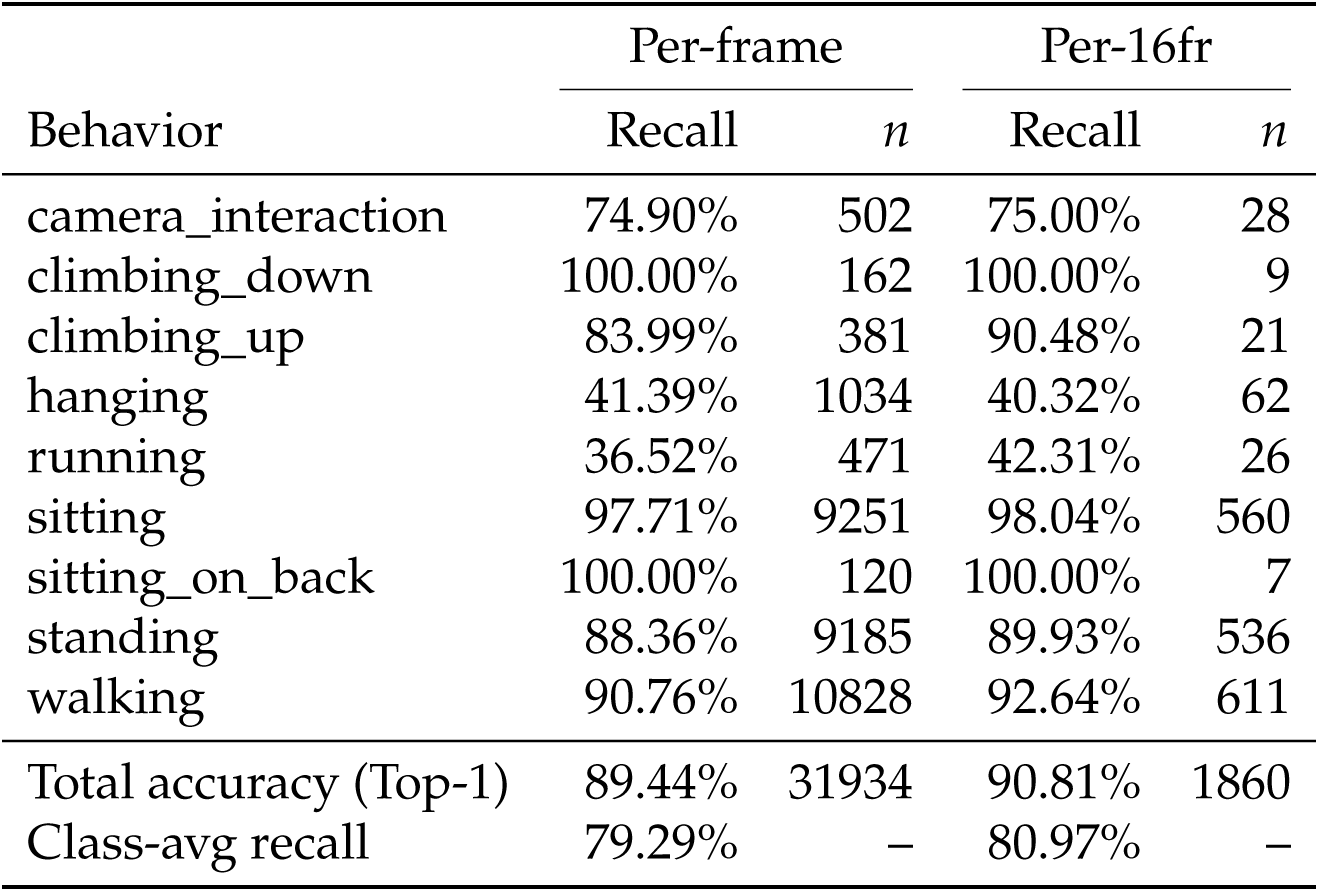
Feral behavior recognition on the PanAf500 test split (75 videos): per-class recall under per-frame vs. per-16-frame-clip evaluation. *n* = number of frames or 16-frame windows.

Additionally, we evaluated FERAL on the PanAf500 dataset under an alternative setup. In this setting, we ignored the bounding box coordinates and used only the behavior labels, aggregating the behaviors of all annotated individuals within a frame into a single multi-label target. For example, if two apes were labeled as *sitting* and one as *climbing*, the resulting frame-level target would be *sitting* and *climbing*. Unlike the methodology described in the original paper, we did not crop the images around individual subjects. As shown in Table 5, performance in this setup is substantially lower, although the model still achieves reasonable results on the most common classes. This reduction in performance is likely due to the weaker supervision signal, as the model must infer behaviors from the entire frame rather than from subject-centered crops. Furthermore, resizing the full frame to 256 × 256 pixels likely removes fine-grained visual details that are better preserved in the cropped-image setting.

**Table 5:** Per-behavior F1 on PanAf500 without cropping. % frames sums to *>*100 because the task is multilabel (a frame can carry multiple behaviors); the *Total* row reports the macro-averaged F1.

| Behavior | Test frames | % frames | F1 |
| --- | --- | --- | --- |
| camera_interaction | 665 | 1.85 | 0.565 |
| climbing_down | 244 | 0.68 | 0.827 |
| climbing_up | 484 | 1.34 | 0.774 |
| hanging | 1504 | 4.18 | 0.465 |
| running | 440 | 1.22 | 0.153 |
| sitting | 11374 | 31.60 | 0.849 |
| sitting_on_back | 439 | 1.22 | 0.000 |
| standing | 10449 | 29.03 | 0.744 |
| walking | 11199 | 31.11 | 0.822 |
| **Macro F1** |  |  | **0.578** |

#### *C. elegans* locomotion

Animals were grown at 20 ^◦^C on nematode growth media (NGM) plates seeded with *Escherichia coli* OP50 bacteria [60]. Experiments were performed on young adult hermaphrodites that were picked as L4s 14–16 h prior to the assay.

The animals express two transgenes: the integrated array *kyIs830* = pSM(F23H12.7p::ReaChR::sl2::GFP) + pSM(myo3p::mCherry) and the extrachromosomal transgene *njEx1552* = pSM(ser-4p::flp, sto3-p::frt::HisCl1::sl2::mCherry, ges-1p::nls-GFP). The animals were not exposed to all-trans-retinol or histamine during the recordings.

Animals were recorded while performing an off-food foraging assay [46, 47], with the change of using a plastic ring (6 mm Clear Mylar Stencil Sheets) instead of CuCl_2_ as a boundary to contain the animals. Preconditioning plates were made the night before by seeding NGM plates with a thin uniform OP50 lawn sixteen hours prior to the start of the assay. 45 minutes prior to the start of the assay, a 1 in × 1 in plastic ring was placed on the preconditioning plate as a boundary and ∼20 young adult worms were picked onto the lawn. 5 minutes before the start of the assay, 8–11 animals were transferred to an unseeded NGM plate to clean off excess food, then transferred to the assay plate, an unseeded 10 cm NGM plate with a plastic ring (2 in × 2 in) boundary to keep animals within the recording field of view. Behavior was recorded for ∼45 min using a Basler ace acA5472-17 *µ*m USB 3.0 monochrome camera at 6 frames per second (fps) and a 3,648 × 3,648 px field of view.

Animals were tracked using custom Python software that segments the worms from background using thresholding and generates tracks of the centroid of segmented worms using intersection over union. This results in tracks including the centroid and mask of the worm.

Behavior features were inferred and classified heuristically using rules adapted from Huang et al. 2006 [49] and Yemini et al. 2013 [48], implemented in Python (code provided), and validated by comparison to human-scored classification (**Extended Figure 2b**).

##### Behavior features

For each frame, it was determined whether the animal was self-intersecting using the area-to-perimeter ratio of the outermost contour of the mask (ratio *>* 3.3). For non self-intersecting frames, masks were skeletonized to obtain the midline of the animal and aligned in the same direction by minimizing the distance between midline points in adjacent frames. The head–tail vector was taken from the first and last midpoints of the animal. Velocity was calculated by taking the difference between centroids four frames (= 0.67 s) apart and dividing by d*t* = 0.67 s. Head versus tail was assigned as the direction along the head–tail axis in which the animal moves more often (the animal moves towards the direction of its head more often than the direction of its tail). Speed was calculated by taking the absolute magnitude of the velocity. Speed was signed by the direction of the velocity relative to the direction of the head–tail vector. *C. elegans* spend significantly more time in the forward state. After following the protocol above, if the number of frames for which signed speed was negative was greater than the number of frames for which signed speed was positive, the sign of the head–tail vector, velocity, and speed was inverted.

##### Behavior classification

Each frame was categorized as one of the motor behavior states: *forwards*, *reversal*, *pause*, or *turn*.

Turning was classified as frames in which either (a) both (i) the midpoint–tail vector (the vector connecting the midpoint and the tail of the worm) was greater in length than the midpoint–head vector (the vector connecting the midpoint to the head of the worm) and (ii) the angle between the midpoint–tail vector and midpoint–head vector was less than 45 degrees, or (b) when the animal was self-intersecting.

Pausing was classified as frames for which |speed| *<* 40 *µ*m s^−1^ and for which the animal was not turning.

Forwards was classified as speed greater than 0 *µ*m s^−1^ and not turning or pausing.

Reversal was classified as speed less than 0 *µ*m s^−1^ and not turning or pausing.

Behaviors were smoothed such that motor states shorter than 0.5 s were classified as the prior state, unless the state began at the start of the track.

### Collective behavior and adult-larva interactions in clonal raider ants (*Ooceraea biroi*)

Invertebrate behavioral research with *O. biroi* was conducted under standard institutional biosafety guidelines.

Stock colonies of *O. biroi* were maintained in constant light at 25 ^◦^C in Tupperware containers (40 × 26 cm) with a ∼2 cm thick plaster of Paris floor. Colonies were fed with frozen fire ant (*Solenopsis invicta*) brood following the lab’s regular feeding schedule (3 times per week) and cleaned and watered once per week, as needed.

For the adult-larva interaction experiments, adult ants and fourth-instar larvae from the same stock colony (clonal line B genetic background) were collected and housed in 5 cm Petri dishes lined with a plaster of Paris floor and kept at 25 ^◦^C. Behavioral assays were performed in a custom-built acrylic chamber with transparent sides and plaster of Paris floor. In each assay, an adult ant was allowed to settle in the chamber for 30 minutes before a larva was introduced. Videos were recorded from the side at 10 frames per second with a FLIR blackfly camera (BFS-U3-50S5C-C) and lens (Computar, MLM3X-MP), using Spinnaker Software Development Kit, at a resolution of 2448 px x 2048 px. Recordings were collected across three consecutive days.

Behavioral annotations of adult ants and larvae were performed manually using ELAN (EUDICO Linguistic Annotator) [61]. We scored adult self-grooming and adult allogrooming of larvae, and recorded the start and end times and duration of these behaviors (in milliseconds).

For the colony tracking experiment, groups of 100 adults and 100 larvae were randomly subsampled from stock colonies and established in Petri dishes (90 x 20mm) lined with a humidified plaster of Paris base. Colonies were allowed to acclimate for four days. Colonies were then continually video recorded at 0.1 fps for 30 days under constant illumination at 25 ^◦^C. Colonies were fed with frozen *S. invicta* brood and cleaned approximately every two days.

Collective raiding events were manually scored from these recordings in BORIS [35] by a single annotator (J.R.), annotating the onset and offset of each synchronized group raid to produce the frame-level ground-truth labels used for training and evaluation.

### Multiclass annotation of raid phases in *Ooceraea biroi*

To assess FERAL on finer-grained collective dynamics than the binary raiding-versus-non-raiding distinction, we analyzed a separate set of *O. biroi* raid recordings acquired in spatially structured arenas, distinct from the colony-tracking dataset above. Colonies on the order of 20 workers were housed in a ∼2 cm-diameter nest connected by a narrow tunnel to a ∼6.5 cm-diameter foraging arena. A prey fire ant (*S. invicta*) pupa was placed in the foraging arena at regular intervals (approximately once daily), and behavior was recorded at 10 Hz. Unlike previously published data, these colonies were not always allowed to settle into the arena before recording, and some were subjected to food deprivation; a subset of these recordings was previously studied in Chandra & Kronauer 2021 [62].

Raids were annotated manually in BORIS [35] by a single annotator (O.S.). Adopting elements of the raid-phase nomenclature previously established for *O. biroi* [58], we annotated each raid into five behavioral states spanning two levels of organization: three collective raid phases — *search*, *recruitment*, and *response* — and two individual ant behaviors — *touching food* and *retrieving food*. The dataset comprised seven video clips from two imaging sessions (five clips used for training and two held out for evaluation), containing six annotated raiding bouts in total. These frame-level state labels were used to train and evaluate FERAL for multiclass prediction.

#### Zebra recordings from drones

Herds of Grevy’s zebras (*Equus grevyi*) were filmed at the Mpala Research Center in Laikipia, Kenya in 2017 and 2018 using DJI Phantom 4 Pro drones (DJI, Shenzhen, China). All fieldwork in Kenya was conducted with the permission of the National Commission for Science, Technology and Innovation and in affiliation with the Kenya Wildlife Service. Drone import and operations were authorized by the Kenya Civil Aviation Authority (KCAA), and data collection protocols were reviewed and approved by Ethikrat, the independent ethics council of the Max Planck Society.

Drone flights were carried out by a licensed pilot assisted by an observer who maintained visual contact with the drone, as well as a ground observer who maintained situational awareness. During filming, the drones were positioned directly above the group at a height of approximately 85 m above ground level and followed the zebra group’s movements. To achieve continuous observations longer than a single drone’s battery duration, two drones were flown in relay: when one drone’s battery became depleted, a second drone was positioned 10 m above the first one. The first drone was then recalled to the launch point, and the second drone was lowered by 10 m to continue following the group.

When groups remained calm and in range, observations covered 3 drone flights, lasting approximately 45–50 minutes in total. During the first two flights, groups were filmed in an undisturbed state. During the third flight, researchers approached the group on foot to elicit a detection and flight response, during which the drone followed the group until they ran out of range. If animals appeared disturbed by the drone (e.g. running away) or moved too far from the launch point, the observation was terminated.

Drone recordings were initially captured at 4K resolution and 60 fps, but were downsampled to 30 fps prior to further processing. A multi-stage pipeline was applied to generate continuous movement trajectories for all animals in the recordings [54]. Recordings were then cropped to generate individual videos of a small square area (160 x 160 or 210 x 210 pixels) centered on each animal, and resized to 256 × 256 pixels before training to match the V-JEPA 2 input shape. Forty-four individual videos from four observations were manually annotated in BORIS [35] to identify bouts of vigilance behavior, defined as the animal standing still with its head raised, and periods when the animal was out of sight, for example due to passing under occluding vegetation. Of these, 35 were used for training, 9 for testing (Extended Table 10).

Initially, a pose estimation approach was used to attempt to identify vigilance bouts. Nine keypoints (nose, head, base of neck, left shoulder, right shoulder, left hip, right hip, base of tail and tip of tail) were annotated in 900 cropped images, and DeepPoseKit [18] was used to train a pose estimation model, which was then applied to the cropped individual-zebra videos (see [18] for full details on pose-model training and performance). To test quantitatively whether vigilance could be recovered from these pose estimates, we trained standard supervised classifiers on the resulting keypoints (**Extended Figure 3a**). For each frame we computed translation- and scale-invariant pose features: keypoint coordinates centered on the confidence-weighted centroid and normalized by their root-mean-square spread, together with keypoint confidences, all pairwise keypoint distances, and per-keypoint angles (72 features). We trained gradient-boosted decision trees (XGBoost [55] and CatBoost [56]; 400 trees, depth 6) to classify each frame as *vigilant* or *other* (out-of-sight frames excluded), using a curated posture dataset of 34,553 frames spanning three observations and 32 individual tracks (class balance 63% *other* / 37% *vigilant*). We evaluated under a leave-one-observation-out cross-validation, in which the model is trained on a subset of recordings and tested on a completely held-out recording, mirroring real deployment. Under this evaluation it collapsed toward chance (balanced accuracy ≈ 0.58 versus a chance level of 0.50; macro-F1 ≈ 0.55; vigilance F1 ≈ 0.40), well below FERAL’s macro-F1 of 0.785 on the same dataset. Both classifiers behaved near-identically.

This failure is consistent with the geometry of the task: vigilance is defined by vertical head elevation, which is poorly captured by the two-dimensional keypoint projection available from the overhead perspective.

#### DeepEthogram

We evaluated FERAL on all six datasets used in the DeepEthogram study. In the original paper, the authors performed five independent train/validation/test splits of the videos (60%/20%/20%), reporting the average performance across the five test sets. Due to the substantially higher computational cost of FERAL, we used a single stratified train/test split per dataset, constructed so that every behavior class is represented in both the training and test partitions, avoiding degenerate splits in which a rare behavior is absent from the small test set. For Mouse-Ventral1 we reused the original DeepEthogram split. For Mouse-Ventral2 and Mouse-Social—the two datasets on which FERAL and DeepEthogram are closest and which are small (16 and 12 videos in total, respectively), making single-split estimates high-variance—we trained on three independent stratified splits and report the best-performing split (Mouse-Ventral2: mean 0.978, range 0.970–0.983; Mouse-Social: mean 0.896, range 0.891–0.900). Checkpoint selection followed the same last-epoch policy used across all datasets (see Model selection).

The six datasets span mouse and fruit-fly behaviors recorded from overhead, ventral, and side views; their species, imaging view, and foreground behavior classes are listed in Table 12, with full acquisition and annotation details given in the original publication [24]. Two features are relevant to the analyses below: Mouse-Social is dyadic—two mice share each frame, so its labels describe relational states rather than a single individual’s posture—and Fly is by far the largest and most temporally fine-grained dataset (*>*3.4 million labeled frames).

#### DeepEthogram: inter-annotator agreement and cross-annotator evaluation

Several DeepEthogram datasets provide labels from multiple independent human annotators (three for Mouse-Ventral1; two for Mouse-Social, Mouse-Homecage, and Mouse-Openfield), which we used to characterize label noise and to test whether FERAL learns the annotator consensus rather than a single annotator’s idiosyncrasies. For each dataset we computed pairwise inter-annotator agreement over all co-annotated frames using raw per-frame agreement, Cohen’sκ, and macro F1 over foreground classes. To relate model error to human disagreement (**Figure 2h**), we partitioned the Mouse-Ventral1 test frames into three buckets by how many of the three annotators agreed on the modal label (all three agree, two of three agree, all disagree) and scored a FERAL model trained on annotator 1 against the modal label within each bucket. In a complementary cross-annotator analysis (**Extended Figure 1d**), we trained a separate FERAL model on each annotator’s labels and evaluated every model against every annotator’s labels on the shared held-out test set, comparing the resulting model-versus-annotator macro F1 to the human-versus-human macro F1.

#### DeepEthogram: annotation-budget analysis

To relate FERAL’s performance to annotation effort, we subsampled training labels at the level of annotated behavioral bouts (contiguous single-behavior segments) on the two most heterogeneous datasets, Mouse-Social and Mouse-Ventral1 (annotator 1). For retention fractions of 75%, 50%, 25%, and 12.5%, we randomly kept that fraction of the bouts of each foreground class and relabeled the frames of the removed bouts as background, leaving the total number of training frames and the test set unchanged, and retrained FERAL from the same initialization with otherwise identical hyperparameters (**Figure 2i**).

#### DeepEthogram: out-of-vocabulary (leave-one-class-out) behavior

To probe how FERAL treats behaviors absent from its training schema, we trained four leave-one-class-out variants on Mouse-Ventral1, each removing one foreground class (face-grooming, body-grooming, digging, or scratching) from the label set and relabeling that class’s frames as background while leaving the remaining classes unchanged. On the held-out test set we measured the F1 of the retained classes and, for the removed class, the row-normalized distribution of the labels FERAL assigned to its frames under the reduced schema (the migration matrix; **Extended Figure 1c**).

### Model Architecture

FERAL fine-tunes the video-understanding backbone to perform frame-level behavioral classification across predefined categories. The model outputs class probabilities for every frame. The complete default configuration used for all reported datasets, including all hyperparameters, is available on GitHub^1^.

#### Backbone evaluation and selection

We systematically evaluated several recent video-understanding architectures as potential encoders for FERAL, focusing on their balance of accuracy, computational efficiency, and ease of deployment. We added an option in the configuration file for users to choose which supported backbone they want to use with FERAL. Currently, three model families (each with multiple distilled versions) are supported: V-JEPA 2, V-JEPA 2.1, and VideoPrism. Our default configuration uses V-JEPA 2. We provide the performance comparison of these models below. **InternVideo2.** InternVideo2 is a 1-billion-parameter transformer model trained in multiple pretraining stages [63]. Despite strong representational capacity, fine-tuning proved prohibitively expensive, requiring up to eight hours for a full run on the CalMS21 dataset using four H100 GPUs. Furthermore, the codebase depended on numerous bespoke modules, complicating installation and development. While performance was promising (91.55% mAP on CalMS21 and 78.2% mAP on the adult-larva ant dataset), these practical limitations rendered InternVideo2 unsuitable for the user-friendly behavioral analysis framework that is the goal of our work.

##### SmolVLM2

We next assessed SmolVLM2, a multimodal video–language model available in configurations ranging from 256 M to 2.2 B parameters [64]. Although integration via Hugging Face Transformers greatly simplified deployment, performance lagged behind V-JEPA 2, achieving only 87.32% mAP on CalMS21. The model processed frames largely in isolation, aggregating temporal information only in a shallow pooling layer. This architectural constraint, compounded by the predominance of non-video modalities during pretraining, limited its ability to capture long-range motion dynamics. Even with extensive regularization (including partial freezing, data augmentations, and label smoothing), SmolVLM2 exhibited overfitting on small datasets and failed to generalize on internal benchmarks, achieving only 47.4% mAP on the adult-larva ant dataset.

##### V-JEPA 2

We chose the publicly released Diving48-fine-tuned V-JEPA 2 ViT-L/16 checkpoint from Meta FAIR as our starting checkpoint rather than the base V-JEPA 2 release, because the Diving48 fine-tuning step was found internally to yield better downstream behavioral classification with no additional cost (`facebook/vjepa2-vitl-fpc32-256-diving48`; ∼300 M encoder parameters; ∼308 M total including the FERAL attention-pool head and classifier) [32]. The encoder was pretrained self-supervised on more than one million hours of internet video and subsequently fine-tuned on the Diving48 human-action dataset [65]. We further fine-tune the upper twelve of twenty-four transformer layers on each FERAL training set. All reported runs use the `transformers` library (version ≥ 4.42) and torch 2.4 with CUDA 12.4. The backbone was pretrained on 32-frame clips at 256×256 resolution; we feed it 64-frame chunks via the model’s positional-embedding interpolation, which we found stable across all benchmarks. Fine-tuning aligns pretrained spatiotemporal representations with the specific requirements of behavioral segmentation, achieving competitive performance across benchmarks with modest data volumes.

##### V-JEPA 2.1

V-JEPA 2.1 shares the architecture of V-JEPA 2 but produces better-aligned spatiotemporal features, and we added support for it following its release. Its default input resolution is 384 × 384, but for consistency with our other experiments we run it at 256 × 256. On our benchmarks it yields similar performance to the V-JEPA 2 backbone.

##### VideoPrism

VideoPrism is a video-understanding model from Google. Because no PyTorch implementation was available, we ported it to PyTorch ourselves^2^. Evaluated on CalMS21, our port slightly exceeded the results reported in the original VideoPrism paper, although it required significantly more compute compared to the frozen backbone methodology. Because V-JEPA 2 achieves better performance after fine-tuning, we retained V-JEPA 2 as the default backbone.

To compare the performance of V-JEPA 2, V-JEPA 2.1, and VideoPrism, we evaluated them on the CalMS21, Adult-larva, and PanAf500 datasets. We used V-JEPA 2 ViT-L (300M parameters), V-JEPA 2.1 ViT-L (300M parameters), and VideoPrism Large (354M parameters). First, we ran experiments using learning rates of 1e-5, 3e-5, 1e-4, 3e-4, 1e-3 on a subset of the Adult-larva dataset. For all models, 1e-4 performed the best, so we ran all models with this learning rate on all three datasets. As shown in **Table 6**, VideoPrism performed worse than the V-JEPA models, and although V-JEPA 2.1 performed slightly better than V-JEPA 2, the difference was not large. Therefore, we decided to keep V-JEPA 2 as our default backbone for now.

**Table 6:**
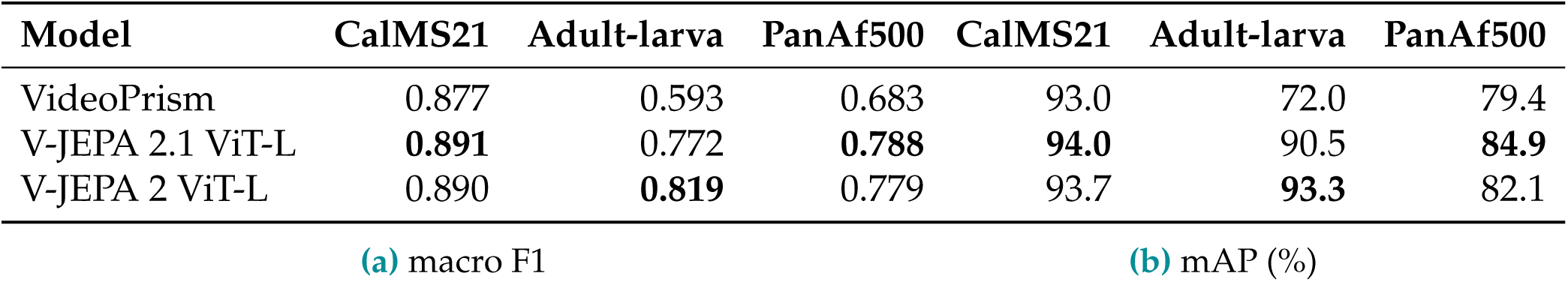
Backbone comparison across datasets. macro F1 (left) and mAP (right) for Video-Prism, V-JEPA 2.1 ViT-L, and V-JEPA 2 ViT-L on CalMS21, Adult-larva ants, and PanAf500, all at LR 10^−4^ on full data.

#### Classification head

To convert spatiotemporal embeddings from the backbone into frame-level behavioral predictions, we designed a lightweight classification head that aggregates contextual information and outputs per-frame logits across behavioral classes.

Each input video chunk is represented as a sequence of thousands of spatiotemporal tokens. These tokens are first enriched with contextual information from the entire sequence by FERAL backbone’s stack of self-attention layers [66]. To map this long sequence onto the temporal resolution of the input video, we employ an attention-based pooling module: sixty-four learnable query tokens cross-attend to the encoder outputs to extract one feature vector per frame.

The resulting pooled embeddings are flattened and passed through a Batch Normalization layer [67], which stabilizes training and controls feature variance, followed by a dropout layer (*?* = 0.5) [68] to reduce overfitting. A final linear projection maps the normalized embeddings to class logits for each frame.

#### Loss function

FERAL employs different loss functions for single-label and multi-label classification. For single label, FERAL is trained using a cross-entropy loss computed at the frame level. To improve generalization and mitigate overconfidence, we apply label smoothing with a factor of 0.1, which encourages the model to distribute probability mass across semantically related classes rather than assigning absolute certainty to a single label.

Because behavioral datasets often exhibit pronounced class imbalance, particularly between dominant “background” states and rare but biologically meaningful actions, we incorporate class-specific weighting into the loss. We found that scaling weights by the square root of the inverse class frequency (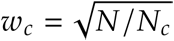, where *N* is the total frame count and *N_c_*the count for class *c*) yielded better performance, amplifying the contribution of underrepresented behaviors without overcompensating, compared with inverse-frequency weighting.

In the multi-label setting, FERAL is trained with binary cross-entropy loss, weighting each class by 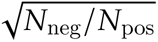 (i.e. the square root of the negative-to-positive ratio) to emphasize rare classes.

We additionally evaluated focal loss (*γ* = 2), which dynamically down-weights easy examples to focus learning on difficult cases, but found no consistent improvement on CalMS21. We therefore decided to use the class-weighted cross-entropy loss.

#### Chunking strategy

Transformer-based video encoders compute pairwise attention across all spatiotemporal tokens, causing computational complexity to scale quadratically with the number of input tokens. As a result, processing entire behavioral recordings in a single pass is infeasible. Consequently, FERAL divides each video into overlapping segments, or *chunks*, of fixed length before processing. Each chunk comprises 64 consecutive frames, resized to a common resolution of 256 × 256 pixels by default.

To capture fine-grained behavioral dynamics, consecutive chunks overlap by 50% (i.e., stride = 32 frames). This design ensures that short behaviors spanning chunk boundaries remain fully visible within at least one receptive field. During training, overlapping windows also increase the effective number of training samples, further improving quality.

During inference, as the model outputs per-frame predictions for each chunk, we employ a unified post-processing pipeline to gather the final frame-level predictions. If a frame received multiple predictions, we averaged the corresponding class probabilities. For frames without a direct prediction, we linearly interpolated probabilities from the two nearest frames with predictions. This approach effectively ensembles the model’s own local predictions.

We benchmarked multiple configurations varying both stride and sampling rate (**Extended Figure 1**) and found that sampling every frame with 50% overlap yielded the best balance between accuracy and training speed.

Note that there is no requirement for the overlap used during inference to match the overlap used during training. Since inference is typically faster than training, a higher overlap can be used at inference time to improve prediction quality without retraining the model. We follow this approach in our *max* configuration, which is trained with 66% overlap (resulting in 3 predictions per frame) and run at inference with 80% overlap (resulting in 5 predictions per frame).

#### Augmentations

To improve generalization across diverse lighting conditions, species, and recording setups, FERAL combines visual (frame-level) augmentations with label-space augmentations. For visual augmentation, we adopted TrivialAugment [69], which, for each image, samples a single transformation uniformly at random from a set of standard image transformations (e.g., brightness, contrast, rotation, color jitter) and applies it at a randomly chosen strength. The same augmentation was applied consistently across all frames within a video, preserving temporal coherence while introducing diversity across video samples.

In addition, we applied MixUp regularization at the batch level [70]. Each augmented sample was formed as a convex combination of two videos, *X* and *X̃*, and their corresponding label sequences, *y* and *ỹ*:

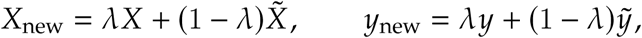

where *λ* ∼ Beta(0.8, 0.8), drawn once per video chunk and applied identically to all frames within the chunk. Because FERAL operates at frame resolution, label mixing was performed element-wise across the temporal dimension. This strategy helps to mitigate overfitting on small datasets.

To test the effectiveness of our augmentation pipeline, we ran evaluations using the default configuration either without MixUp augmentations or without TrivialAugment on CalMS21. As shown in Table 7, augmentations appear to provide a performance boost, with MixUp having a slightly larger effect than TrivialAugment.

**Table 7:** Ablation of augmentation strategies on CalMS21.

| Run | MixUp | TrivialAugment | F1 | ΔF1 | mAP (%) | $\Delta$ |
| --- | --- | --- | --- | --- | --- | --- |
| Original | ✓ | ✓ | 0.883 | – | 94.5 | – |
| No TrivialAugment | ✓ | – | 0.890 | +0.008 | 94.5 | –0.1 |
| No MixUp | – | ✓ | 0.881 | –0.002 | 94.3 | –0.3 |
| No augmentations | – | – | 0.880 | –0.002 | 94.1 | –0.4 |

#### Training

We use the AdamW optimizer [71] with weight decay 0.1, peak learning rate 4 × 10^−5^, a linear warm-up over the first 20% of iterations followed by cosine decay, and batch size 4. Models are trained for 10 epochs, which we found sufficient for convergence across benchmarks.

Unlike the approach used in the VideoPrism paper [33], which freezes the backbone and trains only shallow classifiers, FERAL fine-tunes the upper twelve of twenty-four transformer layers of V-JEPA 2, aligning high-level spatiotemporal embeddings with behavioral structure. To further support generalization and out-of-distribution performance, we allow users to enable an optional weight-averaging feature. When activated, FERAL maintains an exponential moving average (EMA) *θ*_EMA_ of the model weights *θ* [72, 73], which at step *t* is computed as:

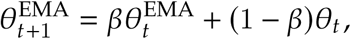

with *β* = 0.999. During evaluation, both the standard and EMA-weighted models are assessed and users can select the best performing option for their application. In our reported experiments, we do not report metrics from EMA checkpoints, as we preferred to keep the evaluation protocol simple rather than risk over-optimizing on them. Because we used *θ* = 0.999, corresponding to updating only 1/1000 of the EMA weights at each step, we observed that the EMA model required at least 1500 training steps to fully warm up. Shorter runs often failed to converge and exhibited substantially worse EMA performance. For our internal experiments that had more than 1500 steps, the EMA checkpoint achieved higher mAP in 54.5% of runs, on average improving by 0.0032 (*n* = 233). Similarly, 55.8% of runs achieved a higher F1 score with the EMA checkpoint, on average improving by 0.0038 (*n* = 197). Despite these gains, the improvements were relatively small and only observed after a sufficiently long warm-up period. Consequently, we disabled EMA in all predefined training configurations, except for *max*.

#### Prediction smoothing

Behavior labels typically do not change rapidly between consecutive frames. If a model frequently flickers between predictions for a behavior, these transitions are often spurious. To address this, we allow users to specify a smoothing window parameter, which averages the predictions for each frame over a temporal window centered on that frame. Prediction smoothing is disabled in the default configuration, but enabled in the *max* configuration with a window size of 9.

#### Presets

There is always a trade-off between quality and speed. To accommodate different application requirements, we provide three FERAL configurations: *lite*, *default*, and *max*. The *default* configuration corresponds to the setting used throughout this paper. The *lite* configuration uses the V-JEPA 2.1 80M backbone, providing approximately a 1.8× increase in training speed while maintaining strong performance. The *max* configuration is similar to the *default* configuration, but increases the temporal overlap during training to 66%, runs inference with 80% overlap, and applies prediction smoothing using a window size of 9. These improvements come at the cost of increased computational requirements, resulting in approximately 1.5× longer training times and 2.5× longer inference times compared to the *default* configuration. Together, these configurations allow users to select an appropriate balance between computational efficiency and predictive quality.

To better illustrate these trade-offs, we evaluated the *lite*, *default*, and *max* configurations on the CalMS21, PanAf500, and Adult-larva datasets. The results are presented in Table 8. We observe that the *max* configuration yields slightly better performance, whereas the *lite* configuration remains competitive while being significantly more efficient.

**Table 8:**
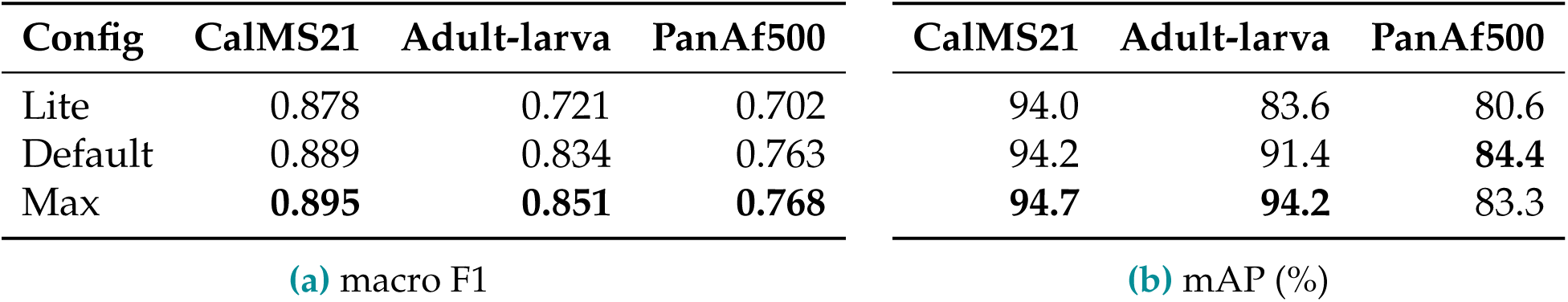
Feral configuration trade-offs across datasets. macro F1 (left) and mAP (right) for the *lite*, *default*, and *max* configurations on CalMS21, Adult-larva ants, and PanAf500.

| Config | CalMS21 | Adult-larva | PanAf500 | CalMS21 | Adult-larva | PanAf500 |
| --- | --- | --- | --- | --- | --- | --- |
| Lite | 0.878 | 0.721 | 0.702 | 94.0 | 83.6 | 80.6 |
| Default | 0.889 | 0.834 | 0.763 | 94.2 | 91.4 | **84.4** |
| Max | **0.895** | **0.851** | **0.768** | **94.7** | **94.2** | 83.3 |

(a) macro F1
(b) mAP (%)

#### Public Checkpoints

We are also releasing^3^ the checkpoints obtained by training on the CalMS21, Adult-larva, and PanAf500 datasets. We provide two versions for each dataset: one compatible with the *max* and *default* setups, obtained using the *max* configuration, and one compatible with the *lite* configuration, produced using the *lite* configuration. Users can use these checkpoints for inference right away or as starting points for fine-tuning on their own datasets.

### Experiments

In the following experiment, each data point was obtained from an independent training run. As a result, some of the configurations were trained multiple times using different random seeds. While these runs produced similar results, small variations are expected due to the stochastic nature of model training. Consequently, performance values may differ slightly between data points.

#### Data efficiency

We tested FERAL’s data efficiency by training on smaller CalMS21 subsets and measuring performance drop versus the full-data baseline, while keeping the test set and all hyperparameters the same. We implemented two complementary subsampling schemes:

**(1) Video-level subsampling.** We randomly sampled subsets of training videos at 50% (mAP 92.0%) and 25% (mAP 93.0%) and trained FERAL on these reduced sets. Because individual recordings vary in length and behavioral composition, smaller subsets (*<*25%) produced high variance across runs: some samples did not contain all classes and the total number of frames varied substantially. We therefore do not report video-level results below 25%. The chunk-level subsampling sweep described next is the one reported in Results (**Figure 2g**).
**(2) Chunk-level subsampling.** To probe sample efficiency under more balanced class distributions, we first processed the full training set into ready-to-train chunks and then randomly subsampled from them. This design mitigates class imbalance and enabled evaluation on much smaller training sets, down to 1% of the original data.

Since expanding datasets typically involves annotating additional videos, video-level subsampling best reflects realistic scaling. However, only chunk-level subsampling allows evaluation under extreme reductions without manual video selection. Under both regimes, training on 25% of the data still exceeded the full-data VideoPrism and Competition Top-1 baselines, and other reduced-data settings maintained strong performance. These results indicate that FERAL’s foundation-model backbone and fine-tuning strategy confer substantial sample efficiency, enabling high performance with limited labeled footage. This is especially valuable in behavioral research, where manual annotation is expensive and time-consuming.

Based on these experiments and our own experience using FERAL for other projects, we found that the most practical way to label data is to annotate a few experimental clips that cover all behaviors of interest, aiming to obtain at least 0.5% of frames for the rarest behaviors. We found that having at least a few thousand frames for each behavior is required for good performance, which corresponds to roughly a few minutes of positive examples at 24 fps.

We also found that it is helpful to start with a subset of the training data to iterate quickly and get a better understanding of which behaviors have enough labels and which require more annotation. `feral train` provides a `--subsample X` argument that samples *X*% of frames from the training partition. We also provide a `--subsample_keep_rare_threshold X` argument, which keeps all chunks containing behaviors rarer than *X* and only subsamples frequent behaviors. This makes it possible to use subsampling even in the presence of rare behaviors.

#### Chunking strategies

We systematically benchmarked chunking strategies to balance computational efficiency and quality. Each configuration is defined by two parameters:

- **Frame stride**: the interval between frames within a chunk (e.g., every frame, every second frame). Larger strides expand the effective temporal window but reduce temporal resolution.
- **Chunk overlap**: the proportion of frames shared between consecutive chunks (e.g., 0%, 50%, 75%). At 0% each frame in the video appears in only one chunk, at 50% in two, at 66% in three, etc. Greater overlap increases computation but improves contextual continuity and yields multiple predictions per frame that can be ensembled to enhance quality.

Performance improved monotonically as stride decreased, with the best results at stride 1 (sampling every frame; labeled as “dense” on **Extended Figure 1a**). Adding overlap yielded substantial quality gains, with 66% overlap producing the highest mean average precision (95.0%). Notably, even sparse settings (0% overlap, sampling every fourth frame) exceeded the competition baseline while requiring 8 times fewer steps than our default configuration. To test whether improvements were simply due to more training steps, we matched the total number of optimization steps by training the base configuration (50% overlap, dense sampling; blue bars on **Extended Figure 1a**) for fewer or more epochs. Dense configurations were clearly superior to sampling every other frame, and moderate overlap was beneficial, although returns diminished at higher overlaps. We therefore adopted 50% overlap with full-frame sampling as the default.

#### Freezing strategies

To evaluate the impact of layer freezing on performance, we incrementally froze six-layer blocks of the 24-layer V-JEPA 2 encoder, proceeding from the input forward, using three different datasets that span different levels of complexity and amounts of data: CalMS21, Adult-larva, and PanAf500. Partial freezing (up to 12 layers) slightly improved test quality, consistent with a mild regularization effect. In contrast, freezing the entire encoder markedly reduced performance (mAP 88.4%), similar to the protocol used by VideoPrism. These results indicate that fine-tuning at least the final layers is necessary to align pretrained features with behavioral segmentation tasks. This is especially apparent when considering the results from the comparatively small adult-larva ant dataset in Table 9.

**Table 9:** Effect of encoder layer freezing on test performance (mAP, %). Six-layer blocks of the 24-layer V-JEPA 2 encoder are frozen incrementally from the input forward.

| Frozen layers | CalMS21 | Adult-larva | PanAf500 |
| --- | --- | --- | --- |
| 0 (no freezing) | 94.2 | 91.6 | 75.5 |
| 6 | **94.4** | 91.4 | 74.4 |
| 12 | **94.4** | **93.4** | 75.4 |
| 18 | 93.7 | 86.2 | **77.2** |
| 24 (all frozen) | 88.4 | 50.7 | 70.4 |

### Metrics

Frame-level mean average precision (mAP) is computed with `sklearn.metrics.average_precision_score` on per-class softmax probabilities for single-label datasets and per-class sigmoid probabilities for multi-label datasets. Throughout this manuscript, mAP refers to the PASCAL-VOC convention: per-class average precision averaged across foreground classes (the “other” class is excluded from the macro-average but retained as a negative for the foreground APs). We additionally report macro F1 — the per-class F1 averaged across foreground classes. For multi-label tasks (e.g., MABe22) we binarize predictions using a fixed sigmoid threshold of 0.85 per class. For the single-label setting, we count a frame as a predicted positive when its logit is the highest among all classes, and negative otherwise. We also report accuracy, defined as the proportion of correctly classified frames. For the multi-label setting, we call a frame correctly classified when predictions for all classes match the target. The performance metrics of FERAL across all benchmark datasets using the same default configuration are reported in Table 13.

#### Metric rationale

Accuracy was not used as the primary metric, since behavioral datasets typically exhibit substantial class imbalance. Datasets often include a frequent “other” class, while biologically interesting events are relatively rare (often *<*1% of frames). In such settings, accuracy overestimates performance by rewarding majority-class predictions. We therefore report mAP, which offers a more sensitive measure of per-class discrimination across thresholds, with macro F1, precision, and recall providing a complementary view. mAP is computed on the raw per-class probabilities described above; the macro F1 score is computed after thresholding continuous model outputs into discrete predictions. For all reported class-averaged statistics, the “other” class is excluded, as it denotes the absence of annotated behaviors rather than a discrete category. We also report top-1 accuracy when comparing against datasets and baselines that historically report it (e.g., DeepEthogram [24] and PanAf500 [53]).

#### Model selection

For all runs, we train the model on the full training dataset and then evaluate it on the held-out test dataset. We do not perform checkpoint selection on a validation split (standard vs EMA, and across epochs), because many of the datasets were small enough that splitting the training set into train and validation either harmed performance by making the training set too small or made the validation split uninformative when it was too small and produced noisy results. This was especially challenging for datasets where some behaviors were present in only a few videos, as the validation set needed to include all behaviors of interest to remain informative. Therefore, we decided to consistently select the non-EMA checkpoint from the last epoch.

We acknowledge that this approach may introduce mild test-data leakage, since we select hyperparameters based on test-set performance. However, we use a single set of hyperparameters across all datasets to avoid overoptimizing for any individual dataset.

#### Run tracking

For tracking training logs, we provide users with two options. Users can either keep all logs private, in which case FERAL prints everything to standard output, or enable automatic logging of all metrics to Weights & Biases (W&B, wandb.ai, [74]), which provides real-time visualization and supports reproducibility. Logged outputs include low-level training diagnostics (e.g., loss and learning rate), per-class average precision at both the chunk and frame levels, aggregated mAP and F1, and qualitative behavioral sequences after each validation and test epoch. Each run generates a unique dashboard URL, making it easy to share, compare, and revisit experiments.

### Engineering and deployment

#### Video preprocessing and seekability

Training requires fast access to arbitrary frames in the video. Some common codecs support only sequential reads and do not allow frame-level seeking, so re-encoding is often necessary. Pre-resizing can also help with HD videos, as decoding large videos may be too slow compared with GPU computation speed. FERAL includes a cross-platform utility that converts input videos to seekable formats and resizes them to 256 × 256 px. If source videos are not high resolution and are already seekable, users may skip re-encoding; resizing will then be performed dynamically during training.

#### Hugging Face integration

To ensure modularity and ease of extension, FERAL leverages the Hugging Face `transformers` library [75, 76] for model management. This abstraction simplifies switching between backbones and streamlines installation compared to earlier architectures (e.g., InternVideo2), which relied on bespoke code and complex dependencies.

#### Training visualization and monitoring

FERAL integrates with Weights & Biases (W&B) [74] for experiment tracking, allowing users to monitor training, validation, and test performance. Logs are stored in the cloud, enabling training across multiple servers while maintaining a unified analysis interface. FERAL comes with a community W&B workspace and a public API key in `default_config.yaml` to lower the onboarding burden for users without a W&B account; `a --no-wandb` flag is provided for fully offline operation, and users can connect their own W&B accounts at any time.

#### Deployment and compute requirements

FERAL *default* requires 24GB of VRAM for training, which fits on high-end consumer GPUs. With gradient checkpointing enabled, the default configuration can also fit within 8GB of VRAM, albeit with longer training times. The Lite version has an even smaller VRAM footprint, delivers slightly lower performance than the default model, but provides nearly 2× faster training. For users without access to GPU hardware, FERAL provides step-by-step deployment guides for GPU cloud platforms (e.g., RunPod) as well as a Google Colab notebook, enabling full training runs at low cost and with minimal setup. All necessary dependencies, including CUDA and PyTorch [77, 78], are pre-installed on these instances, allowing users to begin training within minutes. Inference uses the same chunk length and stride as training (64-frame chunks with 50% overlap) and runs faster than real time on all tested GPUs (see Table 3 for a complete GPU comparison).

## AUTHOR CONTRIBUTIONS

P.S. and J.R. conceptualized the study, developed the method, designed experiments, and performed analyses. P.S., J.R., and L.B.V. co-wrote the manuscript. B.R.C., B.K., and I.D.C. provided the zebra dataset, and B.R.C. supported the application of FERAL to the zebra dataset. F.B. provided the *C. elegans* dataset. D.D.F., D.J.C.K., J.Z., O.S., T.K., and V.C. provided the ant datasets (D.D.F. and J.Z.: adult-larva dataset; T.K. and V.C.: ant colony datasets; O.S.: annotations for the non-binary colony dataset classification). The authors listed between B.R.C. and I.D.C. (F.B., V.C., D.D.F., T.K., B.K., O.S., and J.Z.) are listed in alphabetical order by surname, as they each contributed videos and annotations. All authors provided feedback on the manuscript.

## DATA AND CODE AVAILABILITY

All source code, training configurations, and documentation for FERAL are publicly available at getferal.ai and https://github.com/Skovorp/feral under the MIT license, compatible with the upstream V-JEPA 2 backbone license. The website and the repository include example datasets, preprocessing utilities, and instructions for local and cloud-based deployment. Data and Supplementary Videos 1–4 used in this publication are available at the GitHub Data Repository (https://github.com/Skovorp/feral_share_data/tree/main); a versioned snapshot of code and datasets has been deposited at Zenodo.

## ACKNOWLEDGMENTS

We thank the members of the Vosshall Lab and the Data Science Platform at The Rockefeller University, as well as Giulio Formenti, Yohann Chemtob, and Emily B. L. Wright for discussion and comments on the manuscript. We thank Christopher Harvey for sharing the per-frame accuracy values from the DeepEthogram study. We thank Cornelia Bargmann, who supervised F.B. and generously provided guidance and support for this project. This work was supported by the Howard Hughes Medical Institute and graduate fellowships from the Price Family Center for the Social Brain and the Boehringer Ingelheim Fonds (J.R.). This work was supported by the Howard Hughes Medical Institute, the Rockefeller University, and the Jonathan and Maya Nelson Center for Artificial Intelligence (P.S.). We gratefully acknowledge the Data Science Platform (DSP) at The Rockefeller University for access to the DSP Cluster and computing equipment used in this study. This work was supported by the National Institute on Deafness and Other Communication Disorders under award number K99DC021506 to D.D.F. D.J.C.K. and L.B.V. were supported by the HHMI Investigator program. B.R.C. received support from the European Union’s Horizon 2020 research and innovation program under the Marie Sklodowska-Curie grant agreement No. 748549. B.R.C. acknowledges support from the University of Konstanz’s Investment Grant program. B.K., I.D.C, and B.R.C. acknowledge support from the Deutsche Forschungsgemeinschaft (DFG, German Research Foundation) under Germany’s Excellence Strategy—‘Centre for the Advanced Study of Collective Behaviour’ EXC 2117-422037984, DFG project number 462886202, the European Union’s Horizon 2020 Research and Innovation Programme under Marie Skłodowska-Curie Grant 860949, the DFG Gottfried Wilhelm Leibniz Prize 2022 584/22, and the PathFinder European Innovation Council Work Programme 101098722. B.R.C. acknowledges support from NVIDIA Corporation’s Academic Hardware Grant Program. F.B. was supported by a Medical Scientist Training Program grant from the National Institute of General Medical Sciences of the National Institutes of Health under award number T32GM152349 to the Weill Cornell/Rockefeller/Sloan Kettering Tri-Institutional MD-PhD Program and by an F31 Predoctoral Fellowship from the National Institute of Neurological Disorders and Stroke of the National Institutes of Health under award number F31NS132477. O.S. acknowledges support from the Israel Science Foundation (ISF) through the Beresheet Program for Integrating Outstanding Young Researchers (Grant No. 4126/25) and by the Maimonides Fund’s Future Scientists Center.

## Footnotes

1 https://github.com/Skovorp/feral/blob/main/configs/default_vjepa.yaml

2 https://github.com/Skovorp/torch_videoprism

3 https://www.getferal.ai/pretrained_checkpoints/

## EXTENDED FIGURES

**Extended Figure 1.**
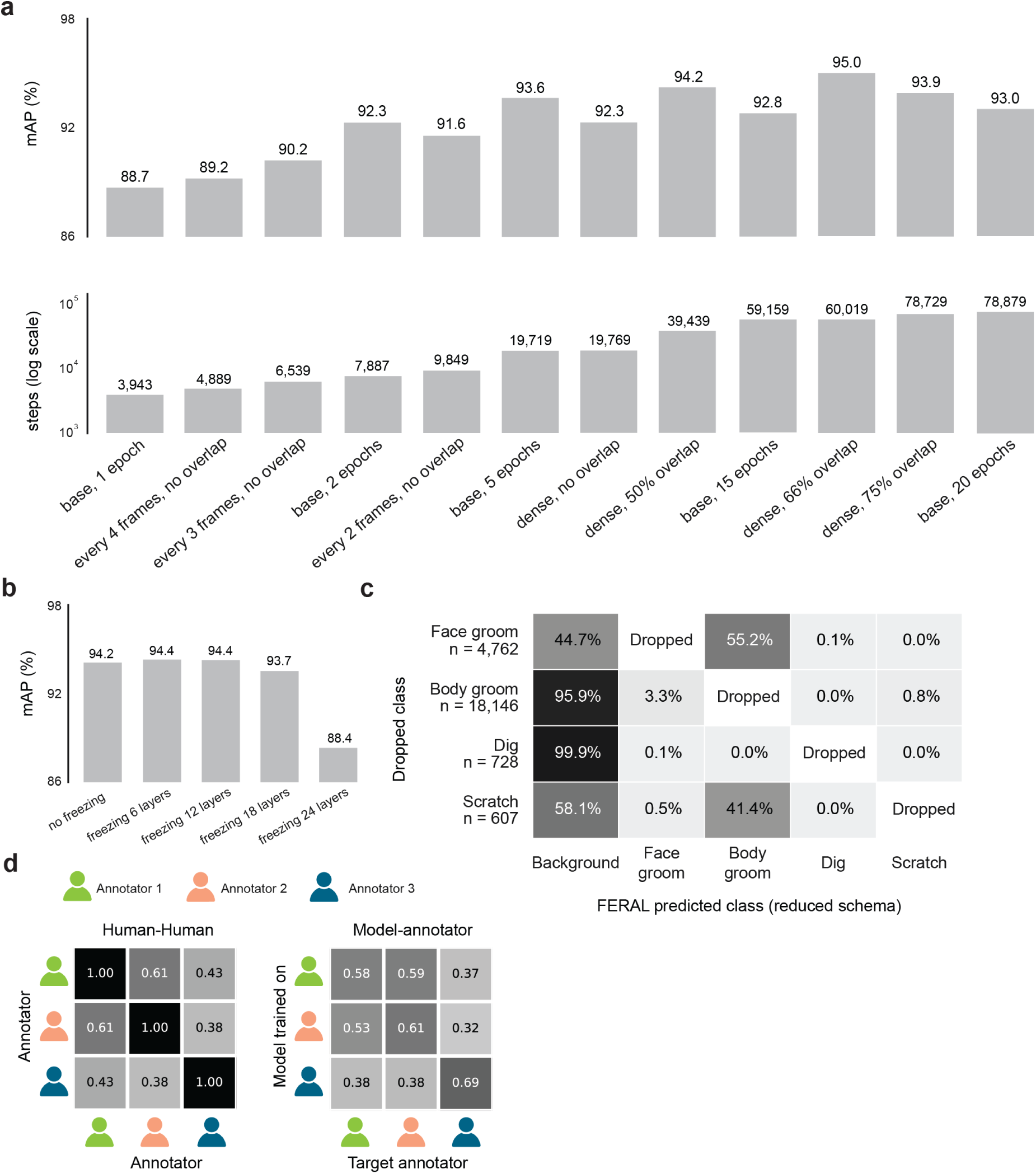
Training-configuration ablations, out-of-vocabulary behavior, and inter-annotator agreement. **(a)** Effect of temporal chunking on the CalMS21 dataset: mean average precision (top) and number of training steps (bottom, log scale) across frame-sampling and chunk-overlap configurations. **(b)** Effect of V-JEPA 2 layer freezing on CalMS21 mAP. **(c)** Out-of-vocabulary (leave-one-class-out) analysis on Mouse-Ventral1: row-normalized distribution of where frames of each held-out class are routed by FERAL under the reduced training schema. **(d)** Inter-annotator agreement on Mouse-Ventral1. Left: pairwise macro-F1 among the three human annotators. Right: macro-F1 between a FERAL model trained on each annotator and each target annotator, showing model–annotator agreement comparable to human–human agreement.

**Extended Figure 2.**
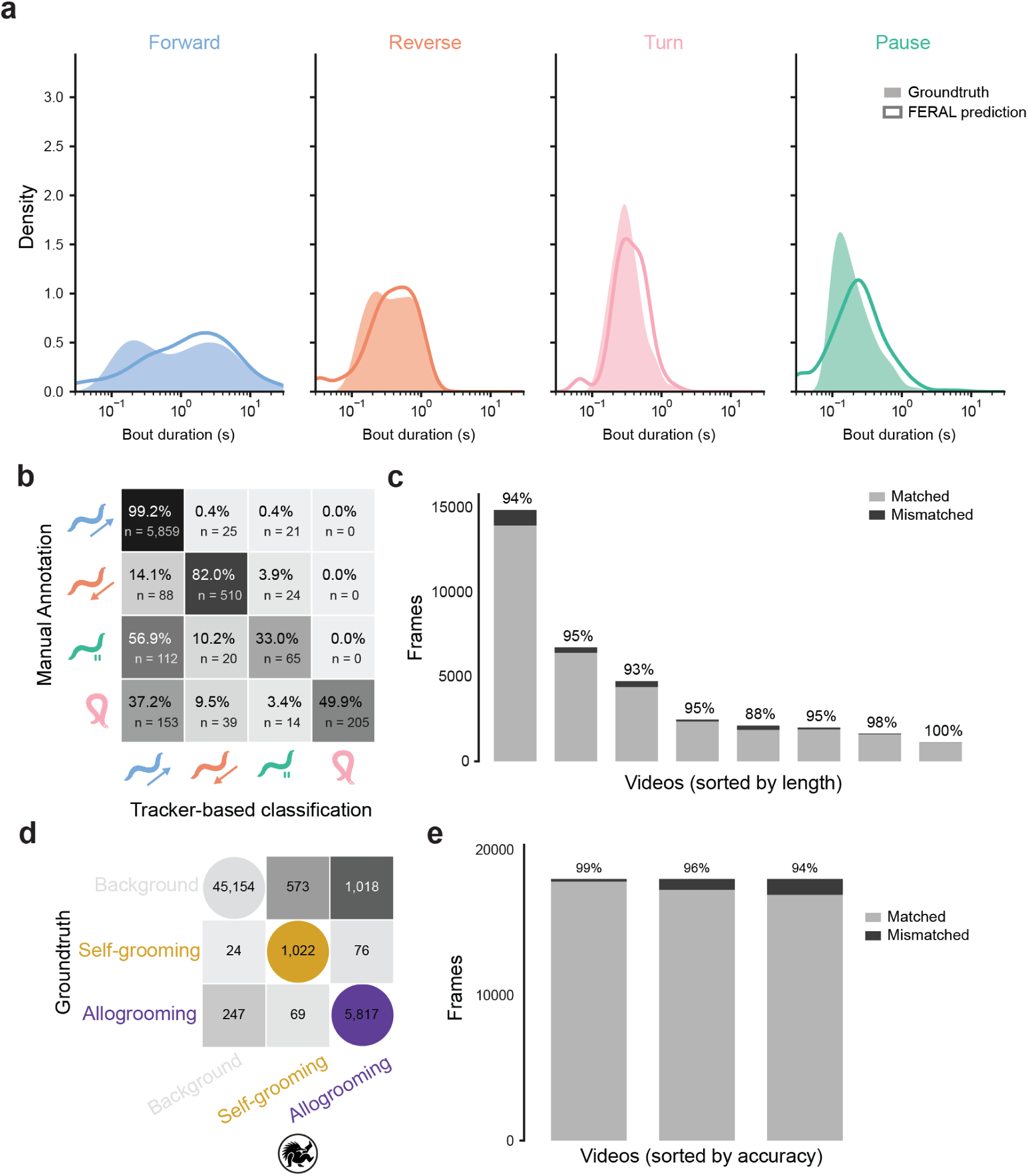
Per-video and confusion-matrix detail for *C. elegans* locomotion and *Ooceraea biroi* adult-larva grooming. **(a)** Distribution of bout durations for the four *C. elegans* locomotor states (*forward*, *reverse*, *turn*, *pause*), comparing ground truth (filled) and FERAL predictions (outline). **(b)** Confusion matrix comparing manual annotation against the tracker-based heuristic classification used as the algorithmic ground truth for *C. elegans*. **(c)** Per-video frame-level accuracy (matched vs mismatched frames) for *C. elegans* videos sorted by length. **(d)** Confusion matrix of FERAL predictions versus expert annotation for *O. biroi* adult-larva interactions (*background*, *self-grooming*, *allogrooming*). **(e)** Per-video frame-level accuracy (matched vs mismatched frames) for adult-larva interactions, videos sorted by length.

**Extended Figure 3.**
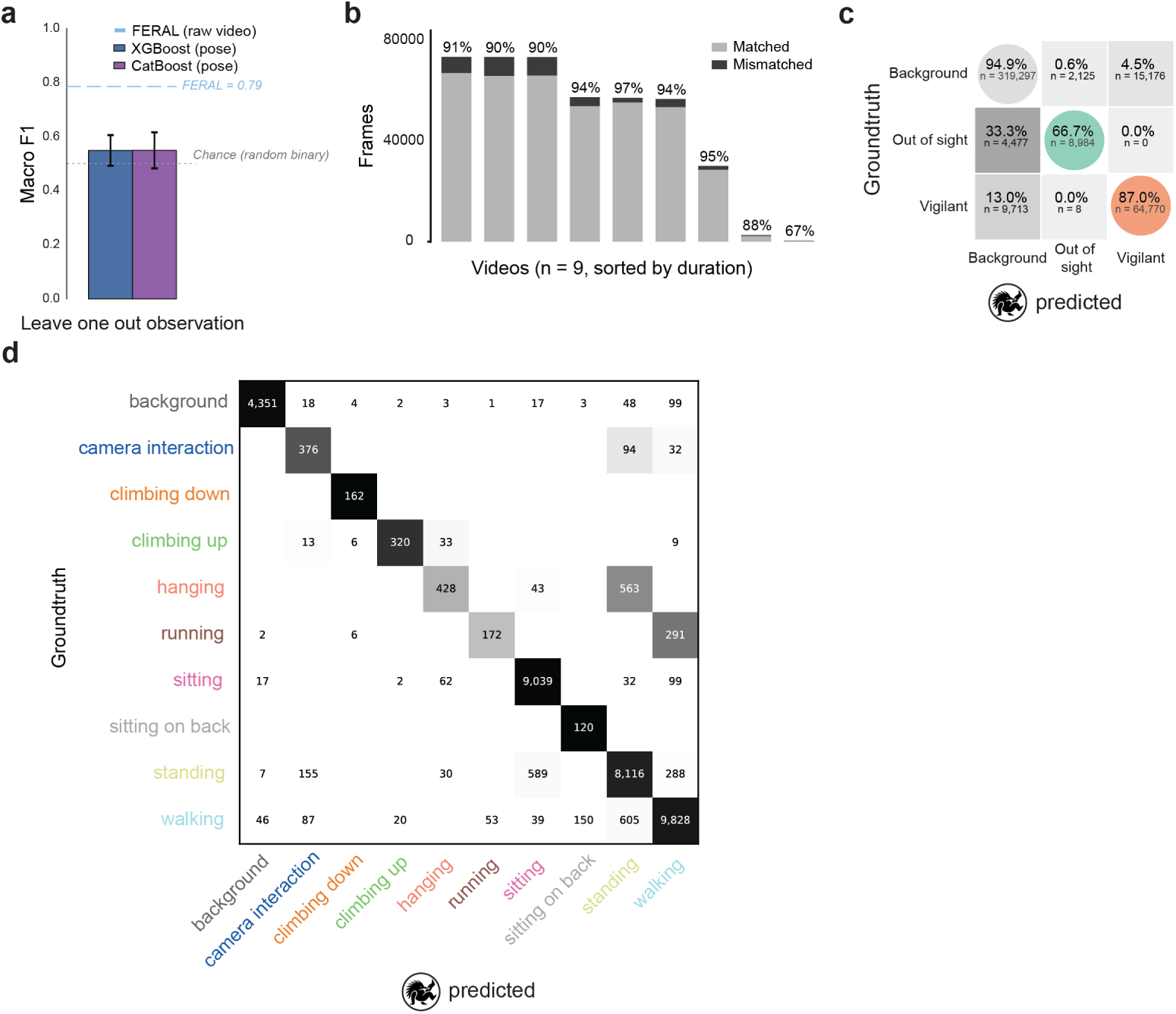
Pose-based classifiers fail on zebra vigilance, and per-video error analysis on the field datasets. **(a)** Vigilant-vs-other classification of zebra frames from DeepPoseKit pose estimates under leave-one-observation-out evaluation: macro-F1 for gradient-boosted classifiers (XGBoost [55], CatBoost [56]) trained on pose features, versus FERAL on raw video (green line, FERAL macro-F1 = 0.785); dashed line, chance for the binary task. **(b)** Per-video frame-level accuracy (matched vs mismatched frames) for the zebra dataset, videos sorted by length. **(c)** Row-normalized confusion matrix of FERAL predictions versus expert annotation for the zebra dataset (*background*, *out of sight*, *vigilant*). **(d)** Confusion matrix of FERAL predictions versus expert annotation for the PanAf500 dataset across the behavioral classes.

## EXTENDED TABLES

**Table 10:**
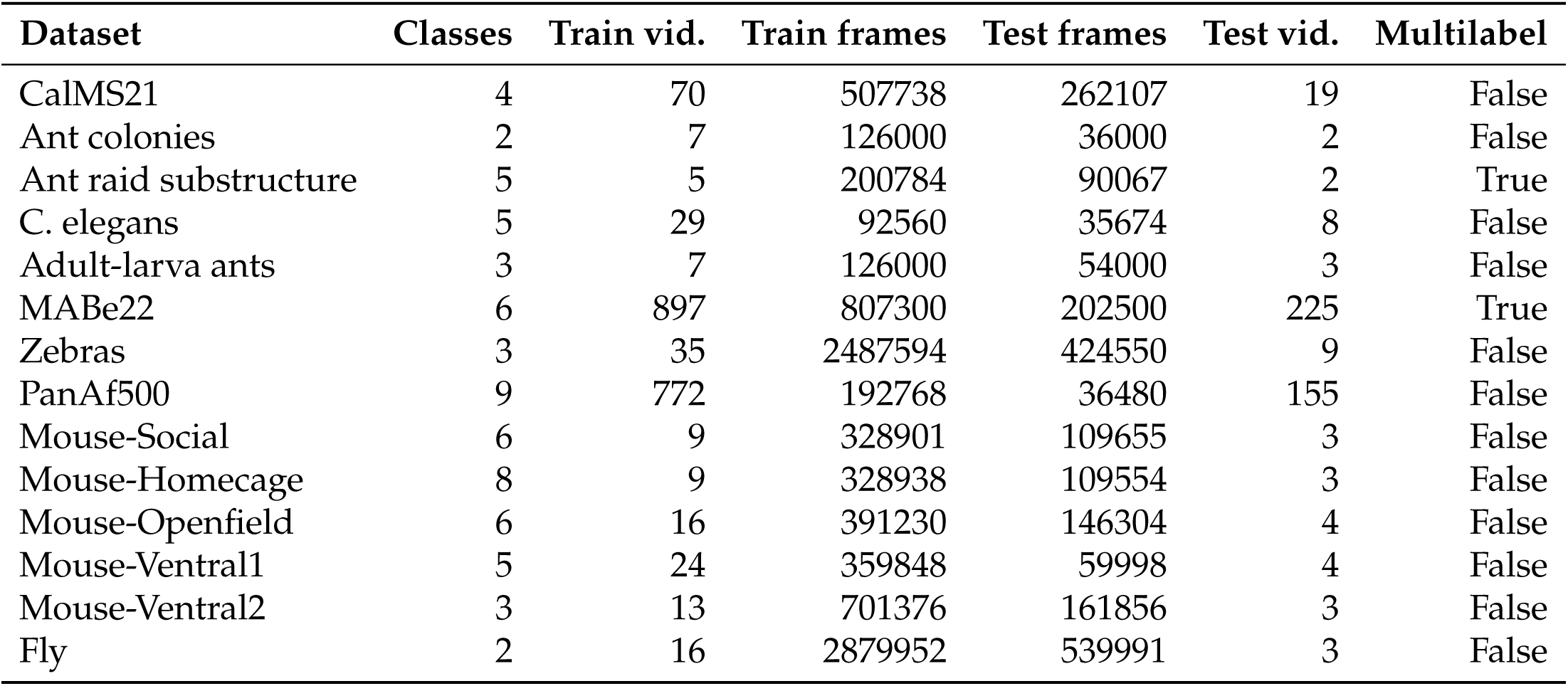
Summary of datasets used in this study. For the PanAf500, C. elegans, and Zebras datasets, we report statistics after cropping each individual into a separate video.

**Table 11:** Label provenance across datasets. How the behavioral labels were generated for each dataset.

| Dataset | Species | How labels were obtained |
| --- | --- | --- |
| CalMS21 | *Mus musculus* (mouse) | Manual expert annotation (challenge-provided) |
| MABe22 (*Sceptobius lativentris*) | Rove beetle | Manual annotation by trained ethologists |
| PanAf500 | Chimpanzee, gorilla | Manual annotation of camera-trap clips |
| *C. elegans* | Nematode | Heuristic / algorithmic — rule-based pipeline from centroid track and midline dynamics (no manual annotation); benchmark of agreement with an algorithmic gold standard |
| Adult-larva ants | *O. biroi* | Manual expert annotation in ELAN (J.Z.) |
| Ant colonies | *O. biroi* | Manual annotation of collective raiding events |
| Zebras | *Equus grevyi* | Manual expert annotation in BORIS (B.R.C.) |
| Mouse-Social | *M. musculus* | Manual annotation (original DeepEthogram authors) |
| Mouse-Homecage | *M. musculus* | Manual annotation (original DeepEthogram authors) |
| Mouse-Openfield | *M. musculus* | Manual annotation (original DeepEthogram authors) |
| Mouse-Ventral1 | *M. musculus* | Manual annotation (original DeepEthogram authors) |
| Mouse-Ventral2 | *M. musculus* | Manual annotation (original DeepEthogram authors) |
| Fly | *D. melanogaster* | Manual annotation (original DeepEthogram authors) |

**Table 12:**
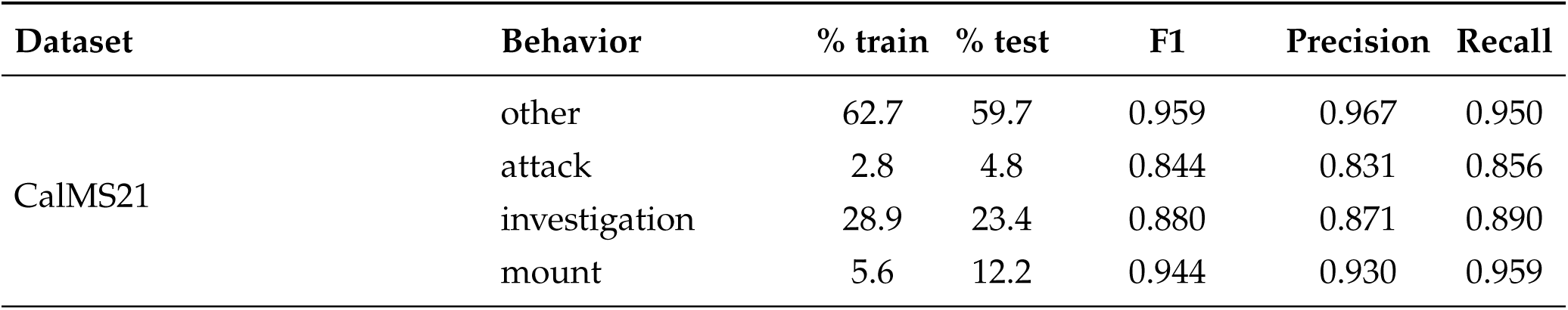
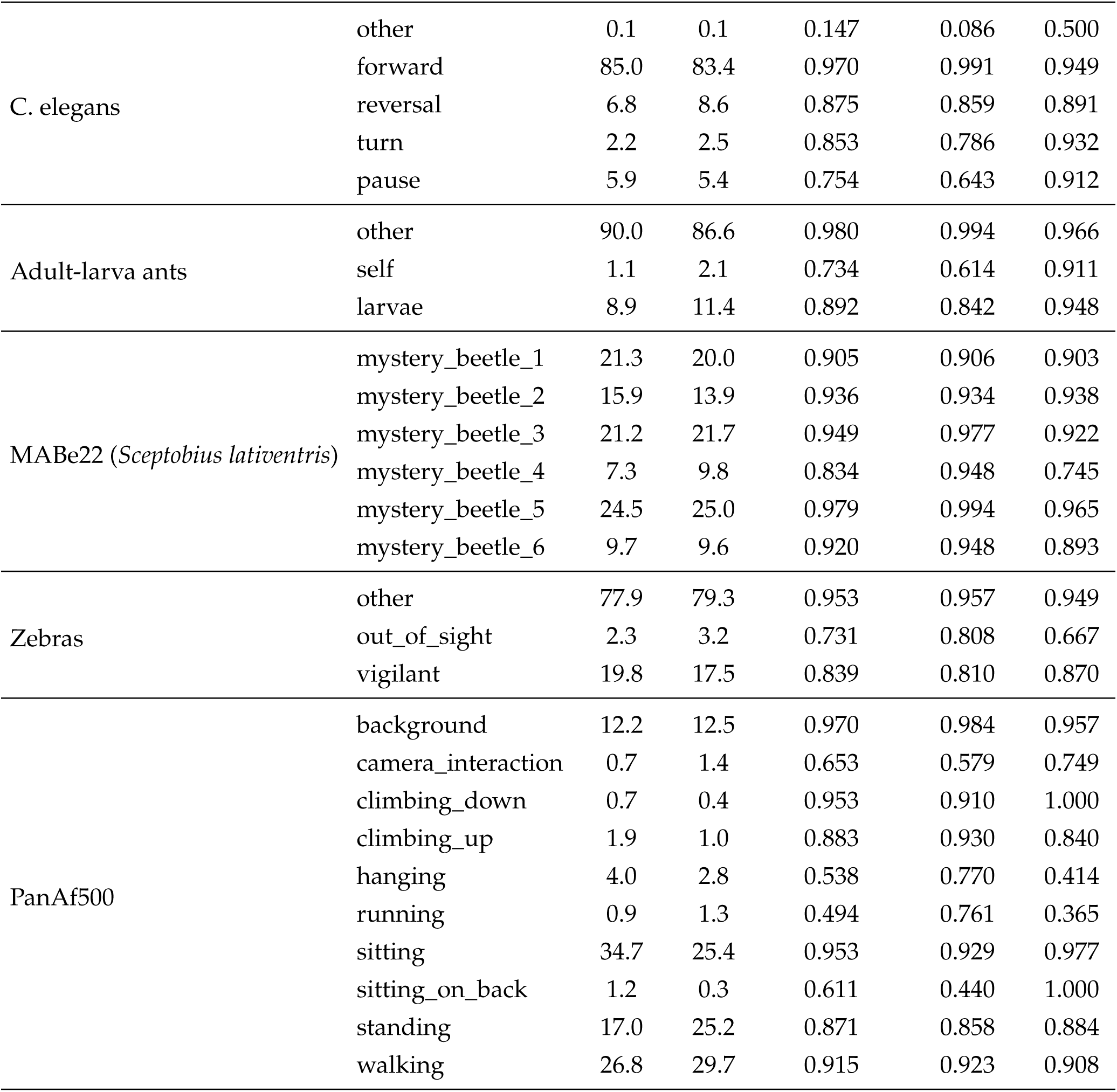
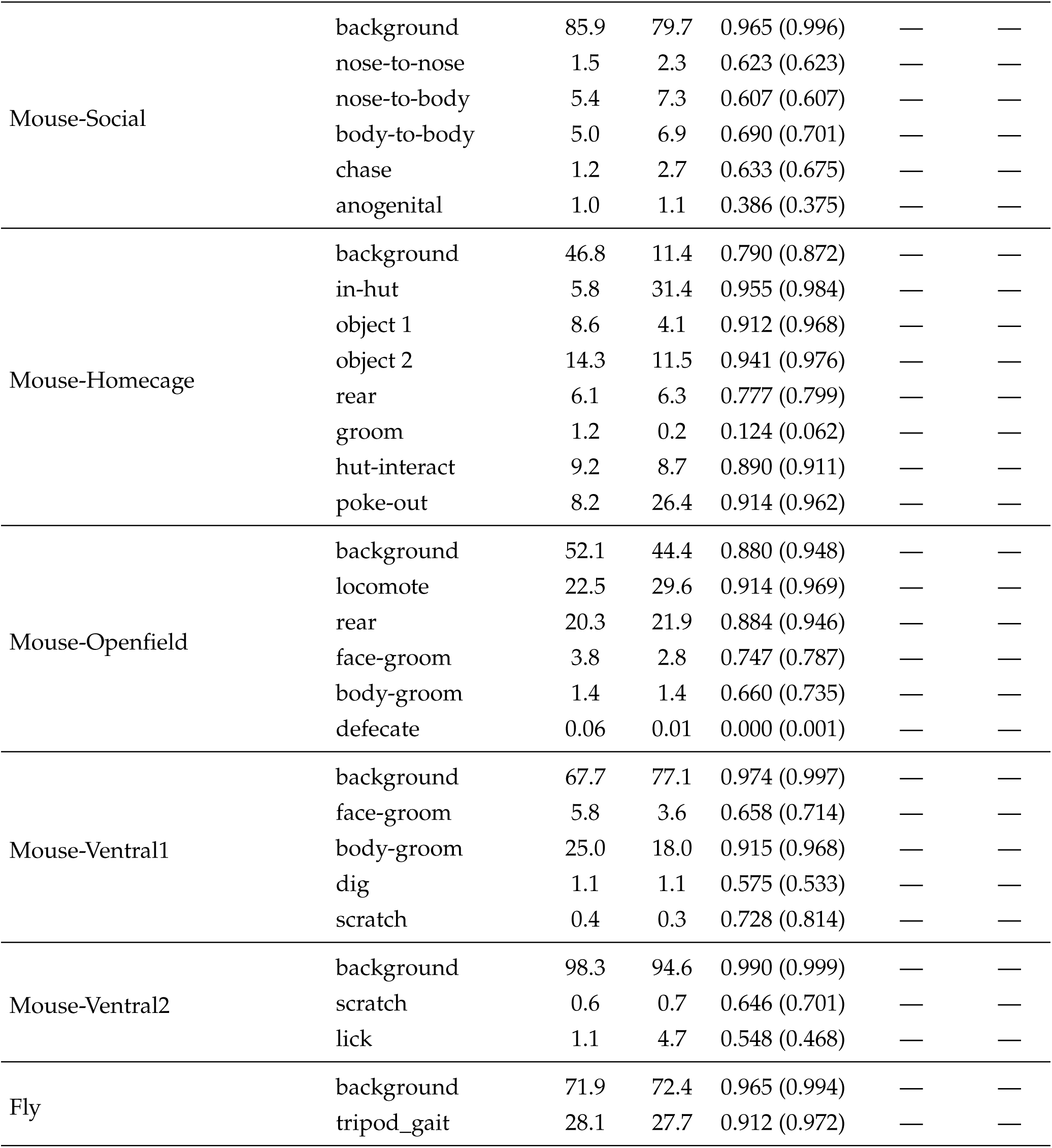
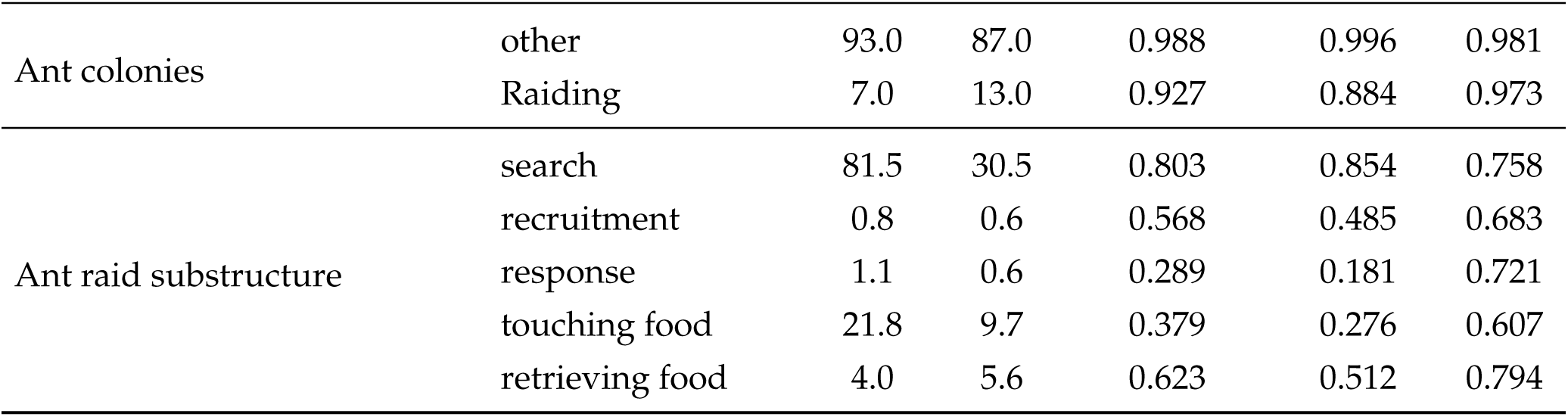
Per-class performance across all evaluated datasets. All experiments used the default configuration. Percentages are the fraction of frames carrying each label in the train / test split; F1, precision, and recall are frame-level on the test split. For the DeepEthogram datasets (Mouse-Social, Mouse-Homecage, Mouse-Openfield, Mouse-Ventral1, Mouse-Ventral2, Fly), per-class precision and recall were not retained; the value in parentheses after the F1 is the per-class average precision (AP).

| Dataset | Behavior | % train | % test | F1 | Precision | Recall |
| --- | --- | --- | --- | --- | --- | --- |
| CalMS21 | other | 62.7 | 59.7 | 0.959 | 0.967 | 0.950 |
|  | attack | 2.8 | 4.8 | 0.844 | 0.831 | 0.856 |
|  | investigation | 28.9 | 23.4 | 0.880 | 0.871 | 0.890 |
|  | mount | 5.6 | 12.2 | 0.944 | 0.930 | 0.959 |
| C. elegans | other | 0.1 | 0.1 | 0.147 | 0.086 | 0.500 |
|  | forward | 85.0 | 83.4 | 0.970 | 0.991 | 0.949 |
|  | reversal | 6.8 | 8.6 | 0.875 | 0.859 | 0.891 |
|  | turn | 2.2 | 2.5 | 0.853 | 0.786 | 0.932 |
|  | pause | 5.9 | 5.4 | 0.754 | 0.643 | 0.912 |
| Adult-larva ants | other | 90.0 | 86.6 | 0.980 | 0.994 | 0.966 |
|  | self | 1.1 | 2.1 | 0.734 | 0.614 | 0.911 |
|  | larvae | 8.9 | 11.4 | 0.892 | 0.842 | 0.948 |
| MABe22 (*Sceptobius lativentris*) | mystery_beetle_1 | 21.3 | 20.0 | 0.905 | 0.906 | 0.903 |
|  | mystery_beetle_2 | 15.9 | 13.9 | 0.936 | 0.934 | 0.938 |
|  | mystery_beetle_3 | 21.2 | 21.7 | 0.949 | 0.977 | 0.922 |
|  | mystery_beetle_4 | 7.3 | 9.8 | 0.834 | 0.948 | 0.745 |
|  | mystery_beetle_5 | 24.5 | 25.0 | 0.979 | 0.994 | 0.965 |
|  | mystery_beetle_6 | 9.7 | 9.6 | 0.920 | 0.948 | 0.893 |
| Zebras | other | 77.9 | 79.3 | 0.953 | 0.957 | 0.949 |
|  | out_of_sight | 2.3 | 3.2 | 0.731 | 0.808 | 0.667 |
|  | vigilant | 19.8 | 17.5 | 0.839 | 0.810 | 0.870 |
| PanAf500 | background | 12.2 | 12.5 | 0.970 | 0.984 | 0.957 |
|  | camera_interaction | 0.7 | 1.4 | 0.653 | 0.579 | 0.749 |
|  | climbing_down | 0.7 | 0.4 | 0.953 | 0.910 | 1.000 |
|  | climbing_up | 1.9 | 1.0 | 0.883 | 0.930 | 0.840 |
|  | hanging | 4.0 | 2.8 | 0.538 | 0.770 | 0.414 |
|  | running | 0.9 | 1.3 | 0.494 | 0.761 | 0.365 |
|  | sitting | 34.7 | 25.4 | 0.953 | 0.929 | 0.977 |
|  | sitting_on_back | 1.2 | 0.3 | 0.611 | 0.440 | 1.000 |
|  | standing | 17.0 | 25.2 | 0.871 | 0.858 | 0.884 |
|  | walking | 26.8 | 29.7 | 0.915 | 0.923 | 0.908 |
| Mouse-Social | background | 85.9 | 79.7 | 0.965 (0.996) | — | — |
|  | nose-to-nose | 1.5 | 2.3 | 0.623 (0.623) | — | — |
|  | nose-to-body | 5.4 | 7.3 | 0.607 (0.607) | — | — |
|  | body-to-body | 5.0 | 6.9 | 0.690 (0.701) | — | — |
|  | chase | 1.2 | 2.7 | 0.633 (0.675) | — | — |
|  | anogenital | 1.0 | 1.1 | 0.386 (0.375) | — | — |
| Mouse-Homecage | background | 46.8 | 11.4 | 0.790 (0.872) | — | — |
|  | in-hut | 5.8 | 31.4 | 0.955 (0.984) | — | — |
|  | object 1 | 8.6 | 4.1 | 0.912 (0.968) | — | — |
|  | object 2 | 14.3 | 11.5 | 0.941 (0.976) | — | — |
|  | rear | 6.1 | 6.3 | 0.777 (0.799) | — | — |
|  | groom | 1.2 | 0.2 | 0.124 (0.062) | — | — |
|  | hut-interact | 9.2 | 8.7 | 0.890 (0.911) | — | — |
|  | poke-out | 8.2 | 26.4 | 0.914 (0.962) | — | — |
| Mouse-Openfield | background | 52.1 | 44.4 | 0.880 (0.948) | — | — |
|  | locomote | 22.5 | 29.6 | 0.914 (0.969) | — | — |
|  | rear | 20.3 | 21.9 | 0.884 (0.946) | — | — |
|  | face-groom | 3.8 | 2.8 | 0.747 (0.787) | — | — |
|  | body-groom | 1.4 | 1.4 | 0.660 (0.735) | — | — |
|  | defecate | 0.06 | 0.01 | 0.000 (0.001) | — | — |
| Mouse-Ventral1 | background | 67.7 | 77.1 | 0.974 (0.997) | — | — |
|  | face-groom | 5.8 | 3.6 | 0.658 (0.714) | — | — |
|  | body-groom | 25.0 | 18.0 | 0.915 (0.968) | — | — |
|  | dig | 1.1 | 1.1 | 0.575 (0.533) | — | — |
|  | scratch | 0.4 | 0.3 | 0.728 (0.814) | — | — |
| Mouse-Ventral2 | background | 98.3 | 94.6 | 0.990 (0.999) | — | — |
|  | scratch | 0.6 | 0.7 | 0.646 (0.701) | — | — |
|  | lick | 1.1 | 4.7 | 0.548 (0.468) | — | — |
| Fly | background | 71.9 | 72.4 | 0.965 (0.994) | — | — |
|  | tripod_gait | 28.1 | 27.7 | 0.912 (0.972) | — | — |
| Ant colonies | other | 93.0 | 87.0 | 0.988 | 0.996 | 0.981 |
|  | Raiding | 7.0 | 13.0 | 0.927 | 0.884 | 0.973 |
| Ant raid substructure | search | 81.5 | 30.5 | 0.803 | 0.854 | 0.758 |
|  | recruitment | 0.8 | 0.6 | 0.568 | 0.485 | 0.683 |
|  | response | 1.1 | 0.6 | 0.289 | 0.181 | 0.721 |
|  | touching food | 21.8 | 9.7 | 0.379 | 0.276 | 0.607 |
|  | retrieving food | 4.0 | 5.6 | 0.623 | 0.512 | 0.794 |

**Table 13:**
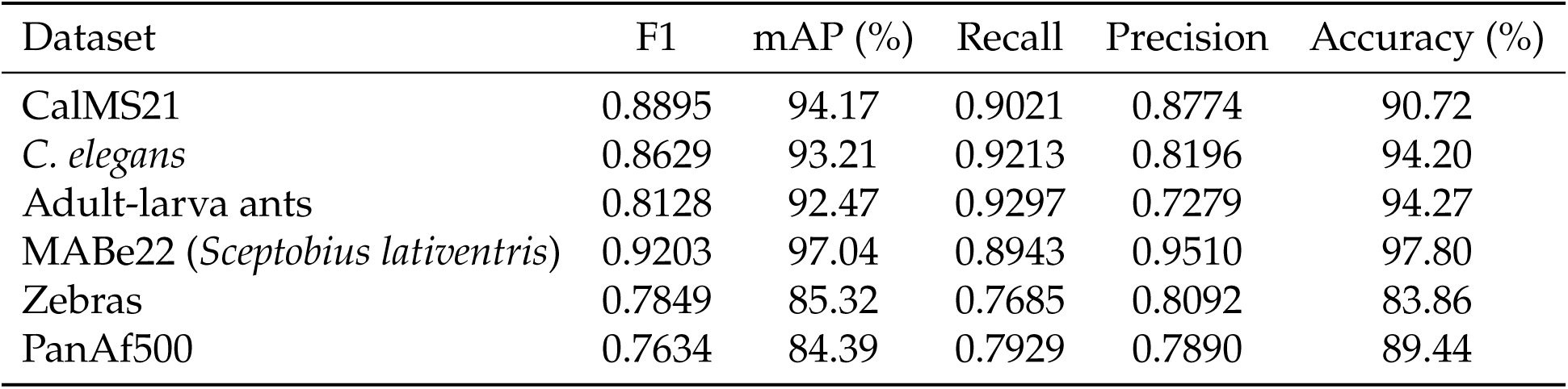
Per-dataset frame-level performance, macro-averaged over non-background behavior classes.

| Dataset | F1 | mAP (%) | Recall | Precision | Accuracy (%) |
| --- | --- | --- | --- | --- | --- |
| CalMS21 | 0.8895 | 94.17 | 0.9021 | 0.8774 | 90.72 |
| *C. elegans* | 0.8629 | 93.21 | 0.9213 | 0.8196 | 94.20 |
| Adult-larva ants | 0.8128 | 92.47 | 0.9297 | 0.7279 | 94.27 |
| MABe22 (*Sceptobius lativentris*) | 0.9203 | 97.04 | 0.8943 | 0.9510 | 97.80 |
| Zebras | 0.7849 | 85.32 | 0.7685 | 0.8092 | 83.86 |
| PanAf500 | 0.7634 | 84.39 | 0.7929 | 0.7890 | 89.44 |

## SUPPLEMENTARY VIDEOS

**Supplementary Video 1 | FERAL on CalMS21 mouse social interactions.** Representative example from the CalMS21 dataset showing raw videos of resident–intruder mouse interactions alongside frame-level FERAL predictions. Link

**Supplementary Video 2 | FERAL segmentation of *C. elegans* locomotor states.** Example recordings of freely moving *C. elegans* with FERAL predictions overlaid on cropped worm-centered views. Link

**Supplementary Video 3 | Adult–larva grooming behavior in clonal raider ants.** Example pair interactions between adult clonal raider ants and larvae, with FERAL predictions overlaid on the raw videos. Link

**Supplementary Video 4 | FERAL on PanAf500 camera-trap recordings of wild apes.** Sample clip from the PanAf500 dataset showing a wild gorilla recorded by a camera trap in its natural habitat. FERAL’s multi-label predictions are displayed frame by frame. Link

